# Heterotrimeric kinesin-2 motor subunit, KLP68D, localises *Drosophila* odour receptor coreceptor in the distal domain of the olfactory cilia

**DOI:** 10.1101/2020.11.24.395905

**Authors:** Swadhin C. Jana, Akanksha Jain, Priya Dutta, Anjusha Singh, Lavanya Adusumilli, Mukul Girotra, Seema Shirolikar, Krishanu Ray

## Abstract

Ciliary localisation of the odour receptor coreceptor (Orco) is essential for insect olfaction. Here, we show that in the *Drosophila* antenna Orco enters the bipartite cilia expressed on the olfactory sensory neurons in two discrete, one-hour intervals after the adult eclosion. Genetic analyses suggest that the heterotrimeric kinesin-2 is essential for Orco transfer from the base into the cilium. Using in vitro pulldown assay, we show that Orco binds to the C-terminal tail domain of KLP68D, the β-subunit of kinesin-2. Reduced Orco enrichment decreases electrophysiological response to odours and loss of olfactory behaviour. Finally, we show that kinesin-2 function is necessary to compact Orco to an approximately four-micron stretch at the distal portion of the ciliary outer-segment bearing singlet microtubule filaments. Altogether, these results highlight an independent, tissue-specific regulation of Orco entry at specific developmental stages and its localisation to a ciliary subdomain by kinesin-2.

**Graphical Abstract:** 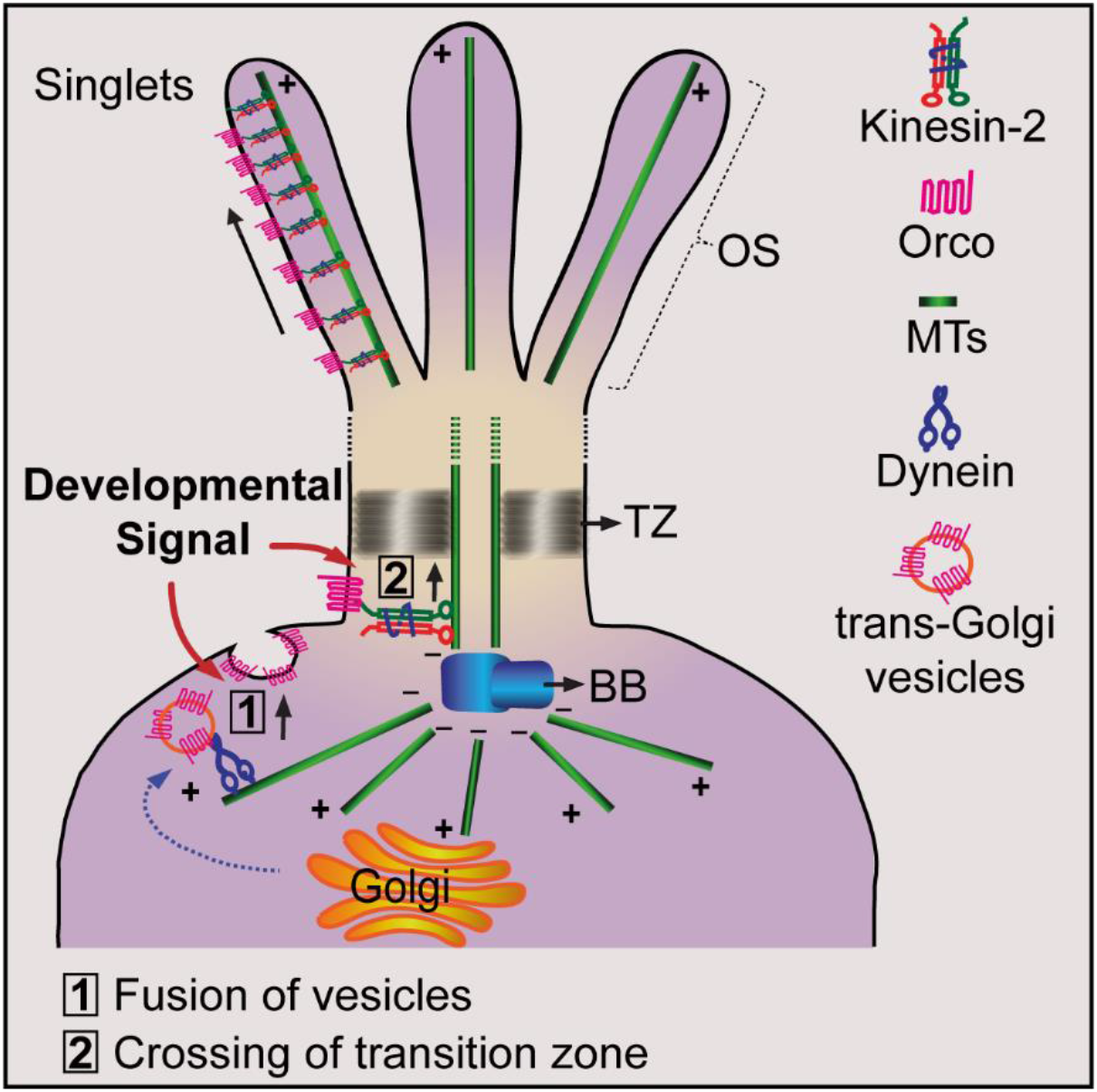

**Author Summary:** Jana, Jain, Dutta et al., show that the odour receptor coreceptor only enters the cilia expressed on olfactory sensory neurons at specified developmental stages requiring heterotrimeric kinesin-2. The motor also helps to localise the coreceptor in a compact, environment-exposed domain at the ciliary outer-segment.

**Highlights:**

- Odorant receptor coreceptor (Orco) selectively enters the olfactory cilia.
- Orco localises in a specific domain at the distal segment of the olfactory cilium.
- Orco/ORx binds to the C-terminal tail domain of the kinesin-2β motor subunit.
- Orco entry across the transition zone and its positioning require Kinesin-2.

## Introduction

Cilia, a microtubule-based sensory appendage, are the cellular antennae that discern the environment and function as a signalling hub (Long and Huang, 2019). The signalling function requires the localisation of transmembrane proteins such as the channel proteins (Hanaoka et al., 2000; Jenkins et al., 2006), G-protein coupled receptors (Omori et al., 2015), growth factor receptors (Gencer et al., 2017) and other membrane components into the ciliary compartment. These proteins localise to the ciliary compartment in response to environmental stimuli and cellular signalling (Kim et al., 2014; Omori et al., 2015). A diffusion barrier isolates both the ciliary membrane and the ciliary cytoplasm from the plasma membrane and the cellular cytoplasm, respectively (Nachury et al., 2010; Vieira et al., 2006). The ciliary compartment is further subdivided into dynamic subcompartments with distinct lipid and protein compositions essential for cell signalling through the cilia (Dyson et al., 2017; Garcia-Gonzalo et al., 2015; Kuhns et al., 2019; Wojtyniak et al., 2013).

Several kinesin family motors are shown to bind to transmembrane receptors and localise them to the cilia. For instance, Kif17, the homodimeric kinesin-2 localises the Dopamine receptor 1 (Leaf and Von Zastrow, 2015), and the Cyclic-nucleotide-gated channel-1 (Jenkins et al., 2006), respectively, into the cilia. Also, interaction with a kinesin-2 motor was indicated to alter the channel functions of Polycistin-2 and Fibrocystin (Wu et al., 2006). In all these cases, the site of kinesin-2 motor function, *i.e.*, whether it helps in crossing the transition zone or afterwards, is unclear. The intraflagellar transport (IFT), driven by the kinesin-2 and cytoplasmic dynein, is proposed to move membrane proteins anterogradely and retrogradely, respectively, inside the cilia (Nachury et al., 2010). Both the IFT and the kinesin-2 family motors are essential for assembly and maintenance of the ciliary cytoskeleton (Evans et al., 2006; Jana et al., 2011; Zhao et al., 2012). Therefore, it was challenging to distinguish the roles of the motor in cilia assembly, compartmentalization and functioning in vivo.

Besides, very little is known about how the membrane-associated proteins move inside the cilium. The issue is best highlighted in our understanding of the Smothend (Smo) transport and localisation process. The Hh stimulated ATPse activity of the kinesin-family motor Costal2/Kif7 is implicated in establishing the Smo signalling compartment at the ciliary tip (He et al., 2014), even though Cos/Kif7 is immotile and has not been shown to bind to either Smo or Gli. Although no direct interaction was shown, β-Arrestin and Kif3A were also implicated in the activity-dependent Smo localisation in the cilia (Kovacs et al., 2008). Independent studies further suggested that signalling-dependent changes of cholesterol level in the ciliary membrane (Weiss et al., 2019), could restrict Smo diffusion along the ciliary membrane after it crosses the transition zone (Ye et al., 2013). This diversity of mechanisms could define the tissue-specific kinetics of hedgehog signalling, but all these were carried out in vitro in tissue-cultured cells. Hence, in vivo experiments are needed to prove the hypothesis.

To address these issues, we investigated the transport of a transmembrane protein, the odour-receptor coreceptor-Orco, into the olfactory sensory cilia in the adult *Drosophila* antenna, which contains multiple classes of sensilla. Each group of these sensilla are innervated with a specific set of olfactory sensory neurons (OSNs) expressing unique odorant receptors (ORs) (Couto et al., 2005). More than 60 different ORs express in these OSNs (Laissue and Vosshall, 2008). Each ORx heterodimerises with Orco forming a functional odorant receptor complex that is essential to localise Orco/ORx to the olfactory cilia (Benton et al., 2006). We have isolated a unique set of kinesin-2 mutants that selectively affect olfactory functions in adult *Drosophila* and established conditional manipulation of kinesin-2 function in the adult olfactory neurons to demonstrate a critical role of heterotrimeric kinesin-2 in Orco localisation into the distal subdomain of an olfactory cilium. Furthermore, we show that Orco selectively associates with the C-terminal tail domain of the kinesin-2 motor subunit, KLP68D, using ex vivo assay. Together, these results indicate that a separate transport paradigm involving heterotrimeric kinesin-2, apart form that invovled in the ciliary assembly, translocates the receptor from the denritic knob across the transitio zone and maintains them at the ciliary outer-segment.

## Results

*Drosophila* antenna contains three morphologically distinct *sensilla* innervated with 2-4 OSNs (Couto et al., 2005; Shanbhag et al., 2000). Bipartite cilia extend from the distal ends of dendrites of the OSNs innervating thee *sensilla* (Figure 1a) and grow into the hollow cuticular shafts (Jana et al., 2011; Shanbhag et al., 2000). Orco expresses in all the OSNs innervating the basiconic and trichoid sensilla and localises in the cilia (Figure 1a; Figure S1). Ectopic overexpression in cholinergic neurons also enriched *Orco:GFP, Or47b:GFP* and *GFP:Or43a* (ORxs) in the cilia inside all types of olfactory sensilla (Figure 1b). The ectopically expressed Or43a and Orco, however, were excluded from the mechanosensory cilia on chordotonal neurons, innervating the Johnston’s organ in the second antennal segment (Figure 1b). Thus, it suggested that Orco/ORx entry is specific to only olfactory sensory cilia, and other molecular systems present in the OSNs facilitate the ciliary access. A similar phenomenon was observed for different G-protein coupled receptors (GPCRs) in *C. elegans* (Wojtyniak et al., 2013).

**Figure 1:**
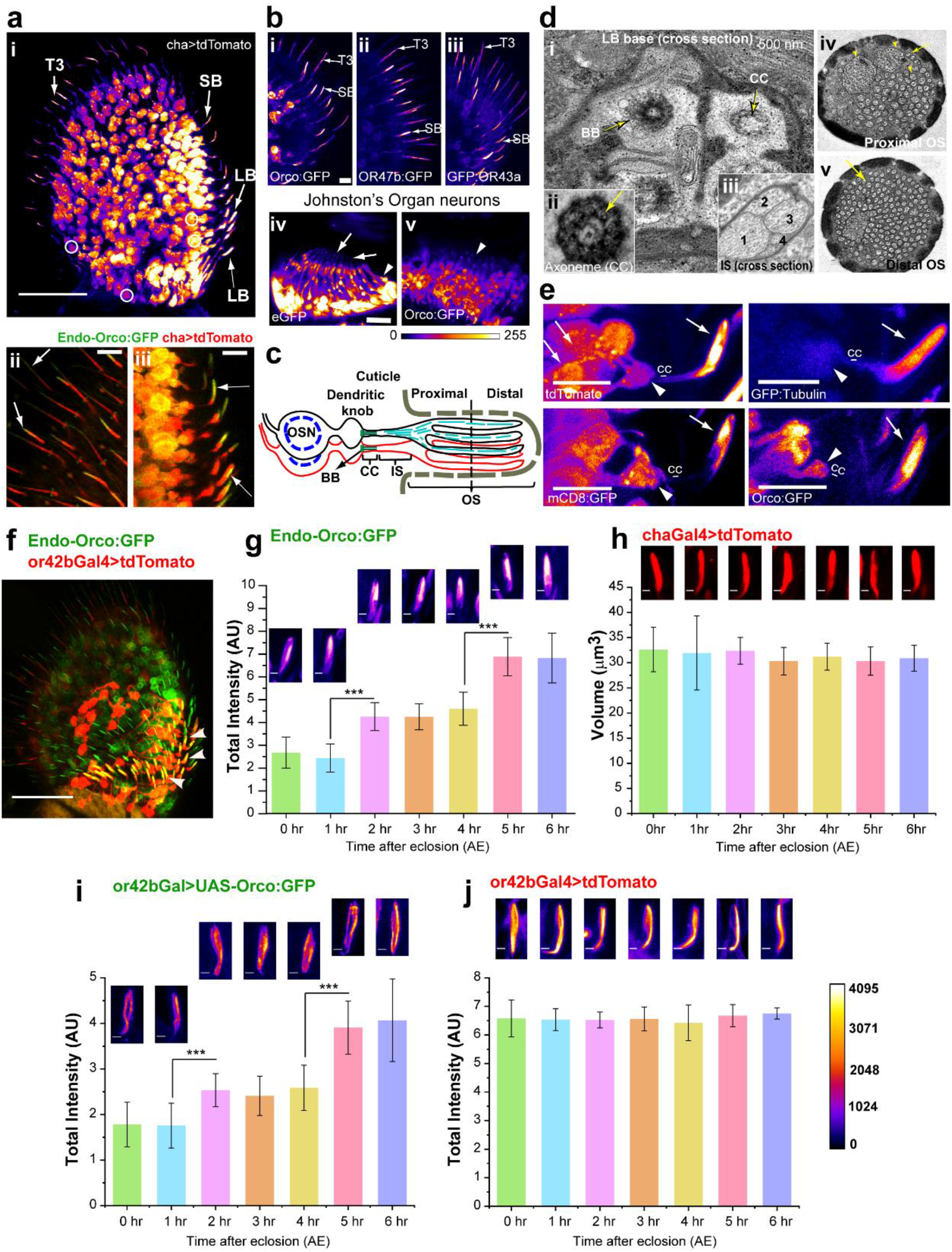
Orco localisation in the olfactory cilia. **a)** Organization of olfactory sensory neurons (OSNs) in the adult third antennal segment marked by tdTomato expression due to *chaGal4*. A few OSN cell bodies are circled white, and the labelled arrows indicate cilia inside small and large *s. basiconica* (SB and LB type) and *s. trichodea* (T3 type) shafts. Lower panels show enlarged regions of large *s. basiconica* (LB) and *s. trichodea* expressing both Endo-Orco:GFP and *cha>tdTomato* (T3). **b)** SB and T3 in the anterior part of the antennae marked by ectopic expression of Orco:GFP **(i)** and Or47b:GFP **(ii)**, and GFP:Or43a **(iii)** by *chaGal4* (arrows). Ectopic expression of soluble EGFP (iv) and Orco:GFP/Or47b:GFP (v), respectively, by *chaGal4* in the chordotonal cilia. Arrowhead marks the cell body of the chordotonal neurons and arrows marks the cilia. **c)** Schematic of an *s. basiconica* innervated by two olfactory sensory neurons (OSNs). **d)** TEM images of transverse sections through the base of an LB-type sensillum show the basal bodies (BB) and the connecting cilium (CC) of four innervating cilia **(i)**. Each CC carries nine doublet microtubule (9+0) bearing axoneme (arrow, **ii**). Sections of the inner (IS, **iii**) and outer segment (OS, **iv**) shows the arrangement of singlet microtubules within a membrane sleeve (yellow arrow). The number of such singlet branches increases at the distal segment. **e)** The tdTomato, Tubulin:GFP, mCD8:GFP and Endo-Orco:GFP localisation in the cilia inside ab1-type *s. basiconica* (arrows-ciliary OS, and arrowheads-ciliary base). **f)** Adult antenna expressing Endo-Orco:GFP and *or42b>tdTomato* marking the ab1-type sensilla (arrowhead). **g, h)** Total fluorescence intensity of Endo-Orco:GFP localised in the ab1-type *s. basiconica* **(g)**, and ciliary volume marked by tdTomato **(h)** expressed in all OSNs from 0-6-h AE. **i, j)** 5xGFP:Orco **(i)** and tdTomato **(j)** enrichment in the or42b cilium inside ab1-type *s. basiconica*. The pairwise significance of difference was estimated using one-way ANOVA test, p-values (*p < 0.05, **p < 0.01, and ***p<0.001) are shown on the plots. Error bars represent as + S.D. Images are shown in a false colour intensity heat map (FIRE, ImageJ®). Scale bars indicate 50 μm **(a-I, ii, iii** and **f i)**10 μm **(b, c)**; and 2 μm **(g, h)**, respectively.

The olfactory cilium extends from a basal body (BB) placed at the distal end of the OSN dendrite (Figure 1c). Each cilium inside the *s. basiconica,* henceforth called basiconic cilia, has a bipartite organisation well-resolved by confocal microscopy. The base consists of a 9+0 arrangement of microtubule doublets (Figure 1d), which supports ~0.5-μm-long connecting cilium (CC) between the ciliary inner-segment (IS) and the basal body (BB). The outer-segment (OS) is extended from the IS and housed within the cuticle shaft. It consists of membranous branches supported by singlet microtubule (MT) filaments (Figure 1d). The OS contains two distinct structural domains. The proximal domain with fewer branches containing multiple singlet microtubule filaments, and highly branched distal domain where each branch has a single microtubule filament tightly invested with the ciliary membrane (Figure 1d). The ab1-type large basiconica are innervated with four OSNs, and one of these expresses OR42b (de Bruyne et al., 2001). We used these sensilla for all quantitative analysis in this study, because of its characteristic features and high levels of Orco enrichment (Figure 1e, f).

### The bulk of the Orco enters cilia after eclosion in two, hourly intervals

Basiconic cilia fully develop by 90-hours after pupa formation (APF) (Jana et al., 2011). Although each ORx expresses at a distinct developmental stage in designated OSNs (Clyne et al., 1999), the *Orco* gene expression begins 80-h APF (Larsson et al., 2004); Figure S1a). In combination with ectopic and overexpression experiments, we tracked the Orco localisation in the developing olfactory cilia in the flies expressing Orco:GFP from its endogenous promoter using an *Orco^fosmid^* stock (Sarov et al., 2016), hereafter, referred to as Endo-Orco:GFP. Although the entire ciliary volume and membrane were marked by soluble GFP and mCD8:GFP (Lee et al., 1999) respectively, during the pupal stages (84-h APF), the overexpression of Orco:GFP and Or47b:GFP using the *chaGal4* driver (expressed from the mid-pupa stage in OSNs) restricted the localisation of Orco and Or47b fusion proteins in the cell body until 96-h APF (Figure S1b). The result was consistent with anti-Orco immunostaining analysis (Figure S1a). We also found that the Endo-Orco:GFP failed to localise to the cilia until 90-h APF. Subsequently, an increasing number of basiconic cilia were marked by the Endo-Orco:GFP in a graded manner (Figure S1c-d), indicating that the Orco entry is correlated to the developmental maturation of the olfactory cilia.

The Endo-Orco:GFP levels attained the equilibrium values within 6 hours after eclosion (AE) (Figure S1e-f), indicating that bulk of Orco/OR enters the cilia within a short period immediately after the adult emerges from the pupal case. Using the ab1-type sensilla marked by the *or42bGal4>tdTomato* expression in the *Orco^fosmid^* background (Figure 1f), we further resolved the entry period to two distinct, less-than-one-hour phases, between 1–2-h and 4–5-h AE, respectively (Figure 1g). The Orco entry into the cilia did not associate with changes in the ciliary volume marked by tdTomato (Figure 1h). Altogether these observations suggested that Orco/OR independently enters the cilia after it is fully developed, and the process is tightly gated. It is also important to note that the availability of the protein does not restrict the Orco entry because we observed an identical pattern in the *or42bGal4>UAS-Orco:GFP* background (Figure 1i) where the Orco transgene is ectopically expressed in a single OSN using *or42bGal4* driver. Together, these observations suggested that the Orco entry is restricted till the end of pupal development, and only initiates after the cilia are matured. Also suggesting that the process must be tightly regulated independently of the *orco* gene expression in each OSN.

### Kinesin-2 is independently involved in maintaining odour reception

Kinesin-2 is essential for cilia assembly and maintenance (Scholey, 2013). It is implicated in the ciliary enrichment of CNGA1 channel and opsins (Jenkins et al., 2006; Trivedi et al., 2012) as well as several other transmembrane receptors (Kovacs et al., 2008; Wu et al., 2006). KLP64D (Kinesin-2α) is an essential subunit of *Drosophila* kinesin-2 (Doodhi et al., 2012). Total loss-of-function (null) mutation in *Klp64D* gene disrupts the development of olfactory cilia as well as reduces the odour-evoked potentials from the antenna (Jana et al., 2011). To assess whether the motor is separately engaged in the Orco/ORx localisation and transport in the adult stage, we isolated viable *Klp64D* alleles with selective olfaction defects in the adult stage (Figure 2a). An initial screen for jump response defects in adult flies isolated five homozygous and viable *Klp64D* mutant alleles (*kj353, kj429, kj925, kj1070,* and *kj1072*) with substantial olfactory defects (Figure 2b-c). The larval chemotactic behaviour of these mutants towards ethyl acetate and Butanol (Table S1); the locomotion of the homozygous mutant adults; as well as the ultrastructure of cilia on chordotonal neurons in the Johnston’s organ; were unaffected in the homozygous *Klp64D^kj^* alleles (Figure S3). These observations confirmed that the *Klp64D^kj^* mutations selectively disrupt adult olfaction, fulfilling the central objective of the screen.

**Figure 2:**
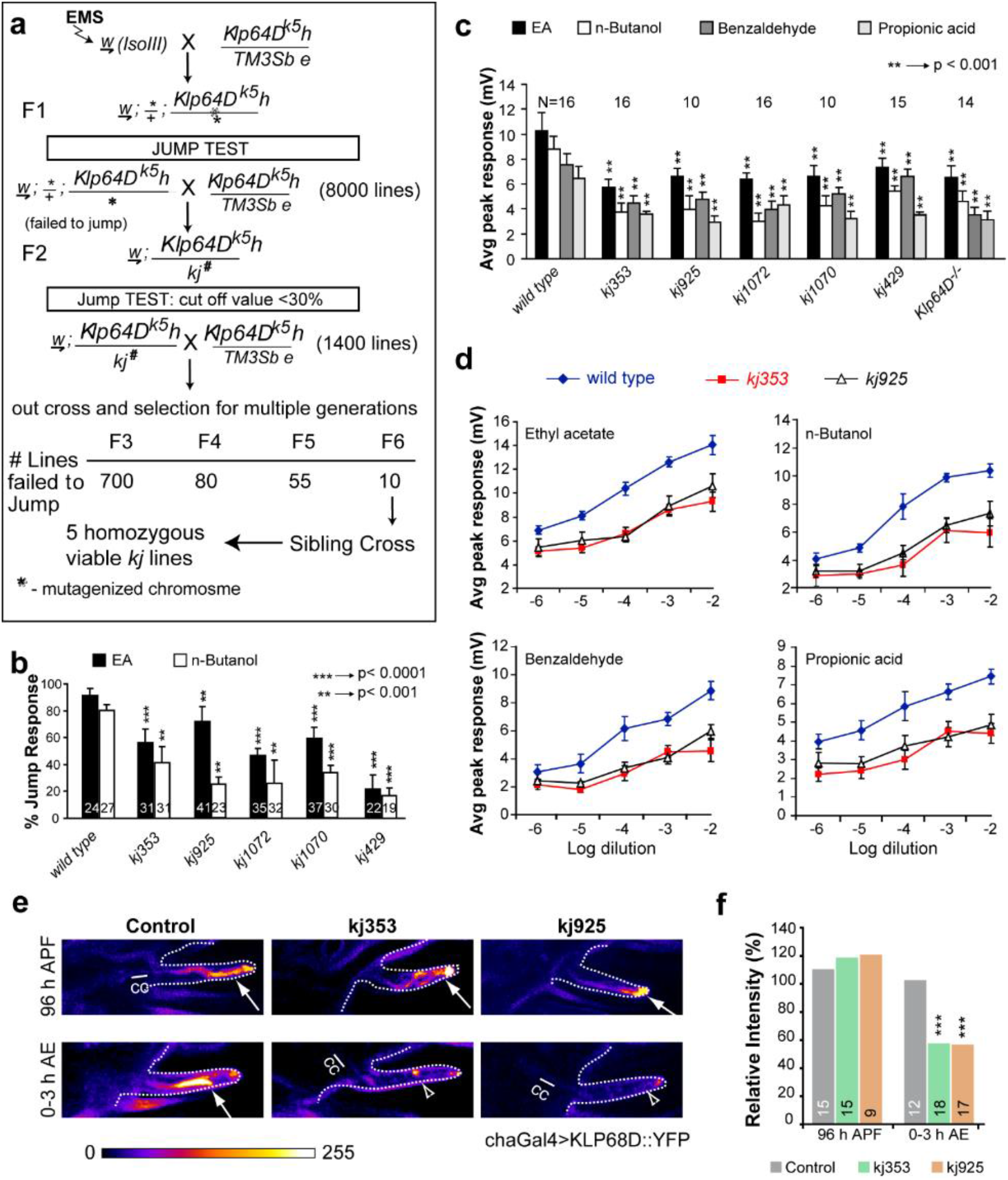
Isolation of homozygous viable *Klp64D* mutant alleles with specific odour response defects in adults. **a)** Flow diagram summarises the ethyl-methane-sulfonate (EMS) mutagenesis screen used for isolating the viable alleles of Klp64D with jump response defect. *Klp64D^k5^* is a homozygous lethal allele. The freshly mutagenised genome was screened for jump response defect mutants in trans-heterozygote combinations over the *Klp64D^k5^* for several generations. Fx (x=1 to 6) indicates the number of generations of outcross with *Klp64D^k5^*. **b)** Jump response indices of wild type (Canton S) and homozygous *kj* isolates. **c)** Electroantennogram (EAG) responses to different odours measured from the homozygous *kj* mutants (see Table 1 and Figure S2 for detail genetic characterisation of the *kj* alleles by EAG). The pairwise significance of difference was estimated using two-tailed Student’s T-test, p-values (*p < 0.05, **p < 0.01, and ***p<0.001) are shown on the plots. **d)** EAG response analyses at different dilutions of four different odorants across the spectrum show that mutations in the homozygous *kj* flies lower the odour sensitivity at the antenna. N ≥10 or as depicted on each set of bars. **e)** KLP68D:YFP localisation in the cilia (expressed by chaGal4) in control, and homozygous *kj353* and *kj925* backgrounds, at 96 hours APF and 0-3 hours AE, respectively. Connecting cilia (CC), and the outer segments (OS, arrows) are shown in each panel (Scale, 10 μm). Images are shown using a false-colour intensity heat map (FIRE, ImageJ^®^). **f)** Relative intensity values of the KLP68D:YFP fluorescence in the cilia estimated against the control group at each stage.

The olfactory sensory responses and the odour-induced neuronal activity are measured using the jump assay and the electroantennogram (EAG), respectively. The aberrant odour-evoked jump responses (Figure 2b) and EAG defects (Figure 2c) displayed by the homozygous mutant flies were recessive in nature. We mapped them to the *Klp64D* locus using genetic non-complementation and transgenic rescue analyses (Table 1). The olfactory defect of the mutants could be rescued by the ectopic expression of *Klp64D* transgene and not by the *Klp68D* (kinesin-2β orthologue) expression (Figure S2a-c). Together, these established that mutations in all the *kj* alleles belong to the *Klp64D* locus. Further, the EAG responses to wide-ranging odours were reduced in the homozygous *Kp64D^kj^* mutants (Figure 2c-d). Of these, ethyl acetate (EA), nButanol (nBut), and benzaldehyde are detected by OSNs innervating *s. basiconica,* and propionic acid by OSNs innervating *s. coleloconica* in the adult antenna (de Bruyne et al., 2001; Hallem et al., 2006). These results indicated that the KLP64D function is necessary for maintaining odour reception by OSNs. A constant downshift of the response voltage at different odorant concentrations further specified the holomorphic nature of the phenotype (Figure 2d).

In the *Klp64D* null alleles, the level of KLP68D:YFP is significantly reduced in the cilia (Jana et al., 2011) because the absence of KLP64D motor subunit destabilises the heterotrimeric kinesin-2 assembly (Doodhi et al., 2012). Therefore, to understand the nature of the kinesin-2 disruption in *Klp64D^kj^* alleles, we estimated the KLP68D:YFP levels in the cilia in wild-type and homozygous *Klp64D^kj353^* and *Klp64D^kj925^* backgrounds, respectively, in the pupae and the adult stages. The ciliary levels of KLP68D:YFP in the mutants was comparable to that of the wild-type at 96 hours APF, but it was significantly reduced in the freshly eclosed (0-3 hours AE) homozygous mutant adults (Figure 2e-f). An in increase in cuticle autofluorescence mostly contributed to the values estimated in the mutants in the adult stage (Figure 2f). This observation suggested that the mutations selectively reduce kinesin-2 entry into the olfactory cilia in homozygous *Klp64D^kj^* mutants after eclosion (AE).

The bulk of the Orco enters the cilia in the adult stage, and, KIF17, the homodimeric kinesin-2, has been reported to facilitate the localisation of CNGA1 channel protein in the mammalian olfactory cilia (Jenkins et al., 2006). Therefore, we reasoned that the loss of kinesin-2 entry into the cilia could affect the Orco/OR transport resulting in the loss of odour reception. Kinesin-2 also transports tubulin into the olfactory cilia (Girotra et al., 2017), and powers the IFT (Scholey, 2013). Both are essential for maintaining the cytoskeletal integrity in the cilia. Hence, disruption of kinesin-2 function could disrupt the ciliary cytoskeleton causing the EAG defect as reported earlier (Jana et al., 2011).

### Loss of kinesin-2 entry reduced the Orco enrichment in cilia

To distinguish between these two possibilities, we compared the enrichment of Orco:GFP, GFP:tubulin84B, and a non-specific membrane marker (mCD8:GFP), respectively, in the ciliary OS in wild-type and homozygous *Klp64D^kj^* mutants. Expression of Orco:GFP in the OSNs by *chaGal4* and *orcoGal4* appeared to localise comparable amounts of the fusion protein in the cilia at 96 hours APF (Figure 3a). However, the ciliary levels were significantly reduced in homozygous *Klp64D^kj353^* adults (Figure 3a, and b), and the expression of KLP64D transgene in the *Klp64D^kj353^* mutant background rescued the defect (Figure 3c). Together these observations suggested that the mutations affect the Orco:GFP enrichment in the cilia only at the adult stage. Interestingly, the difference in ectopic expressions did not influence the Orco:GFP levels in the ciliary OS (Figure 3b), indicating that the ciliary entry of Orco is a robust process, developmentally regulated, and independent of its ectopic expression timing.

**Figure 3:**
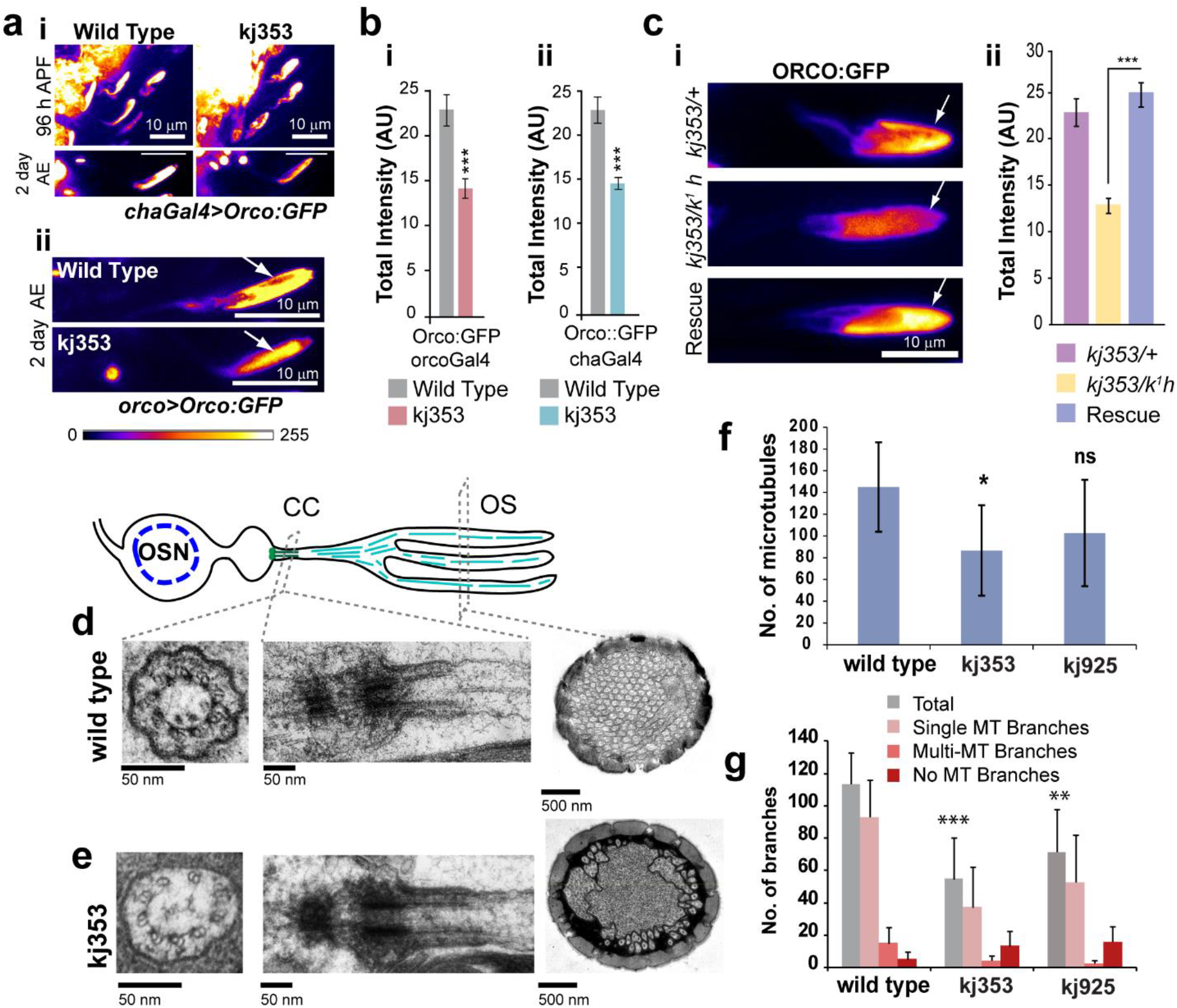
**a-c) Mutations in the *Klp64D* gene reduced Orco localisation in the adult olfactory cilia. a)** Orco:GFP localisation in cilia of *s. basiconica* (arrow), in 96-h APF and two-day-old control and homozygous *Klp64D^kj353^* mutant antennae. The Orco:GFP transgene was separately expressed using *chaGal4* (i) and *orcoGal4* (ii) to test the effects of different levels of recombinant protein expression on its cilia localisation. **b)** Total fluorescence intensity of Orco::GFP driven by *orcoGal4* **(i)** and *chaGal4* **(ii)** in the cilia of *s. basiconica* of two-days-old control and homozygous *Klp64D^kj353^* mutant adults. **c)** Orco:GFP localisation in the ciliary OS of control (*chaGal4>UAS-Orco:GFP/+; Klp64D^kj353^*/+), hemizygous *kj353* (*chaGal4>UAS-Orco:GFP/+; kj353/Klp64D^k1^*) and rescued (*chaGal4>UAS-Orco:GFP/UAS-KLP64D-TevHis*; *Klp64D^kj353^/Klp64D^k1^*) adults. **c)** Total fluorescence intensity measured in the different genetic backgrounds. The pairwise significance of difference was estimated using two-tailed Student’s T-test, p-values (*p < 0.05, **p < 0.01, and ***p<0.001) are shown on the plots. Error bars stand for + S.E.M. All Images are shown in a false colour intensity heat map (FIRE, ImageJ^®^), and scale bars indicate 10 μm. **d-g) Ultrastructure of the cilia inside *s. basiconica* in the wild-type and homozygous *Klp64D^kj^* antennae**. Schematic illustrates the microtubule organisation in cilia of s. basiconica. TEM images of sections through the s. basiconica cilia in one-day-old wild-type (d) and kj353 (e) antennae showed the presence of basal bodies (arrows) and connecting cilia (arrowheads). Panels show transverse and horizontal sections through the BB (arrows) and CC (arrowheads). **f, g)** Histograms describe the total number of singlet MTs **(f)** and the distribution of different types of branches **(g)** found in the OS in wild type and the mutants. The pairwise significance of difference was estimated using one-way ANOVA test, p-values (*p < 0.05, **p < 0.01, and ***p<0.001) are shown on the plots. Error bars represent as + S. D., N > 7, for all sets.

Consistent with this conjecture, we found that the total GFP:tubulin84B level and the ciliary volume were unaltered and there was a marginal increase in the cytoplasmic GFP and mCD8:GFP levels in the cilia in homozygous *Klp64D^kj353^* background (Figure S4). Also, TEM analysis of the antennae of 1-day-old, homozygous, *Klp64D^kj353^* and *Klp64D^k925^* mutant adults failed to identify any significant defect in the ciliary microtubule (Figure 3d-f). Occasionally the structure of the connecting cilium appeared disrupted (Figure 3e), and there was a minor reduction in the number of MTs (Figure 3f). Therefore, we concluded that the kinesin-2 function in the adult stage might not be essential for maintaining the ciliary cytoskeleton. Although the levels of mCD8:GFP were not altered, the number of membrane-bound branches bearing singlet MTs were significantly reduced in the homozygous *Klp64D^kj^* backgrounds (Figure 3g). A combination of negative and positive curvature lipids, such as phosphatidic acid and sphingomyelin, would be required to tightly curve the membrane into a narrow cylindrical shape around the MTs. Also, seven-pass transmembrane-domain receptors like Orco associates with sphingomyelin and cholesterol. Therefore, we reasoned that a loss of Orco transport could result in the loss of associated lipids form the OS, which could reduce the number of branches.

To further test whether kinesin-2 is indeed required for endogenous Orco transport at the adult stage, we sought to knockdown kinesin-2 levels just before the eclosion and study its effect on the endogenous Orco enrichment and compare it with that in the homozygous *Klp64D^kj353^* background. We knocked down kinesin-2 in the adult OSNs by driving Klp64D and Klp68D dsRNAs using *orcoGal4,* which expresses after 80-h APF (Larson et al., 2004) and the RNAi sets in ~16 hours after the dsRNA expression. Therefore, the *orco>KLP64D RNAi* would set in only from 96-h APF, approximately 10 hours before AE. We found a significant reduction of Endo-Orco:GFP in the kinesin-2 RNAi background at 0h AE, and although the level appreciated in the subsequent stages, it remained significantly lower, and delayed as compared to the control (Figure 4b-c). The Endo-Orco:GFP localisation profile in the ciliary OS in *Klp64D^kj353^* homozygous background (Figure 4a) was similar to that of the kinesin-2 RNAi background, suggesting that the kinesin-2 function is indeed required for the progressive increase of Orco localisation into the ciliary OS. Furthermore, the localisation of Jupiter:GFP in the Klp64D RNAi and the Klp68D RNAi backgrounds, respectively, were unaffected (Figure S4d), indicating that the loss of kinesin-2 at the adult stage is unlikely to disrupt the ciliary MT. We also observed that disruption of kinesin-2 function led to Orco accumulation before the transition zone at the dendritic knob (Figure 4), indicating that kinesin-2 motor is required to move Orco across the transition zone.

**Figure 4:**
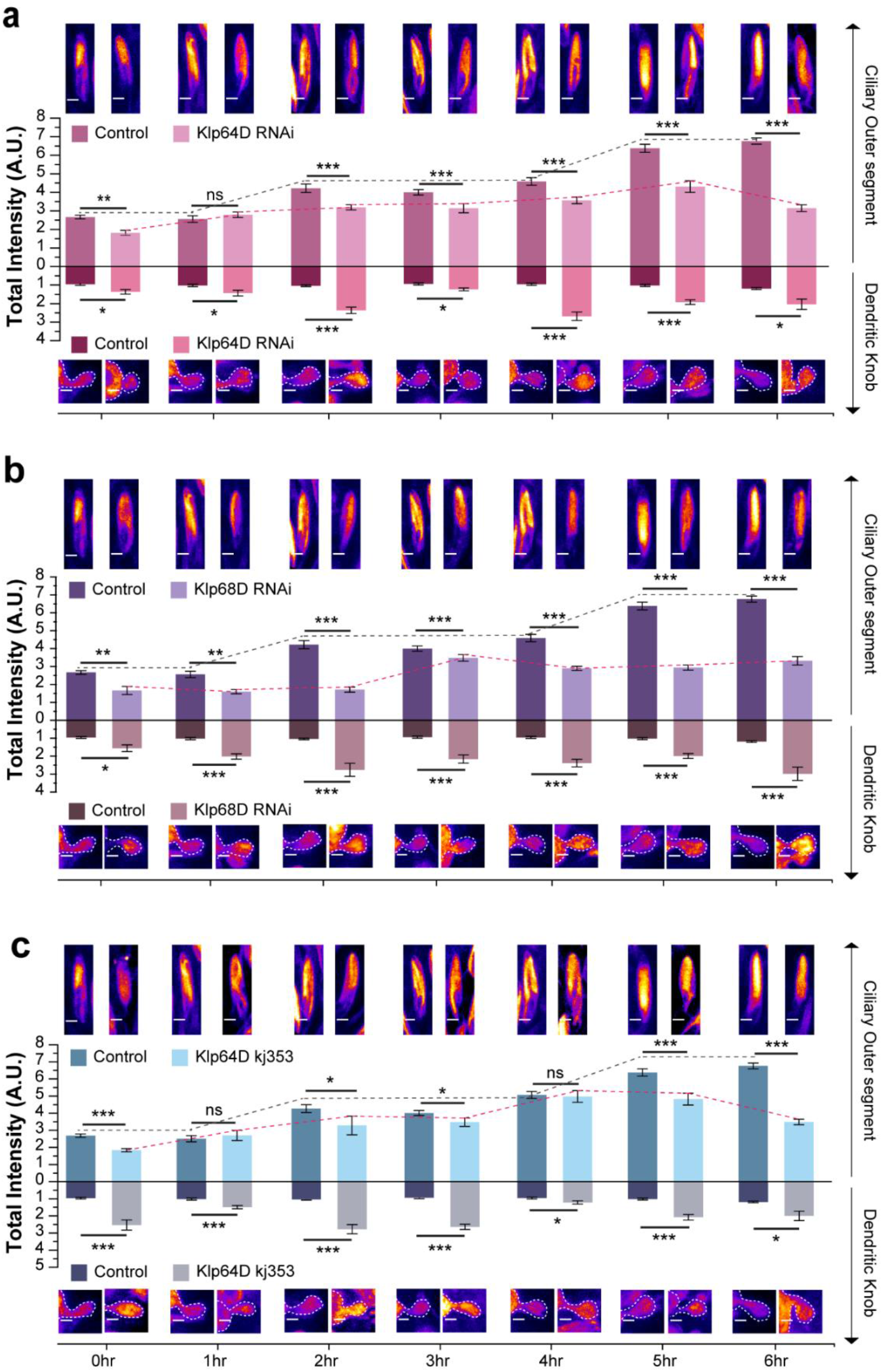
Kinesin-2 is necessary for the entry of Orco in the olfactory ciliary OS. **a-c)** Total fluorescence intensity of Endo-Orco:GFP in the ciliary OS and in the dendritic knob region of OSNs. The pairwise significance of difference was estimated using one-way ANOVA test, p-values (*p < 0.05, **p < 0.01, and ***p<0.001) are shown on the plots. Error bars represent as + S.E.M. Images are shown in a false color intensity heat map (FIRE, ImageJ^®^), and scale bar indicates 2 μm.

### Orco associates with the C-terminal tail domain of the Klp68D (kinesin-2β) motor subunit

Next, to supplement the genetic evidence, we tested whether the heterotrimeric kinesin-2 motor (Figure 5a) would directly engage Orco/ORx using copurification assays. The N-terminal domains of kinesin-2 motor subunits bind to microtubule and generate the force (Scholey, 2013). The coiled-coil stalk domains in the middle are involved in heterodimerisation (Doodhi et al., 2012) and bind to the accessory subunit kinesin associated protein (KAP) (Doodhi et al., 2009) and an IFT complex subunit (Baker et al., 2003). The C-terminal tail domains bind to cargoes like Choline acetyltransferase, Acetylcholinesterase, Rab4, and tubulin (Sadananda et al., 2012; Scholey, 2013). The anti-GFP antibody precipitated both the KLP68D (kinesin-2β) and DmKAP (Figure 5b) from the head extracts of *chaGal4>UAS-Orco:GFP* adults, suggesting that Orco interacts with the heterotrimeric kinesin-2. Further, Orco:GFP was selectively pulled-down using the GST-KLP68D-Tail fragment (Figure 5c) which mapped the potential Orco binding region to the tail domain of KLP68D (2β) motor subunit. In addition, OR47b was also copurified with the KLP68D tail by affinity chromatography of the head extract (Figure 5d). Together, these results suggested that Orco/ORx associates with the heterotrimeric kinesin-2 motor, which subsequently transports it into the cilium.

**Figure 5:**
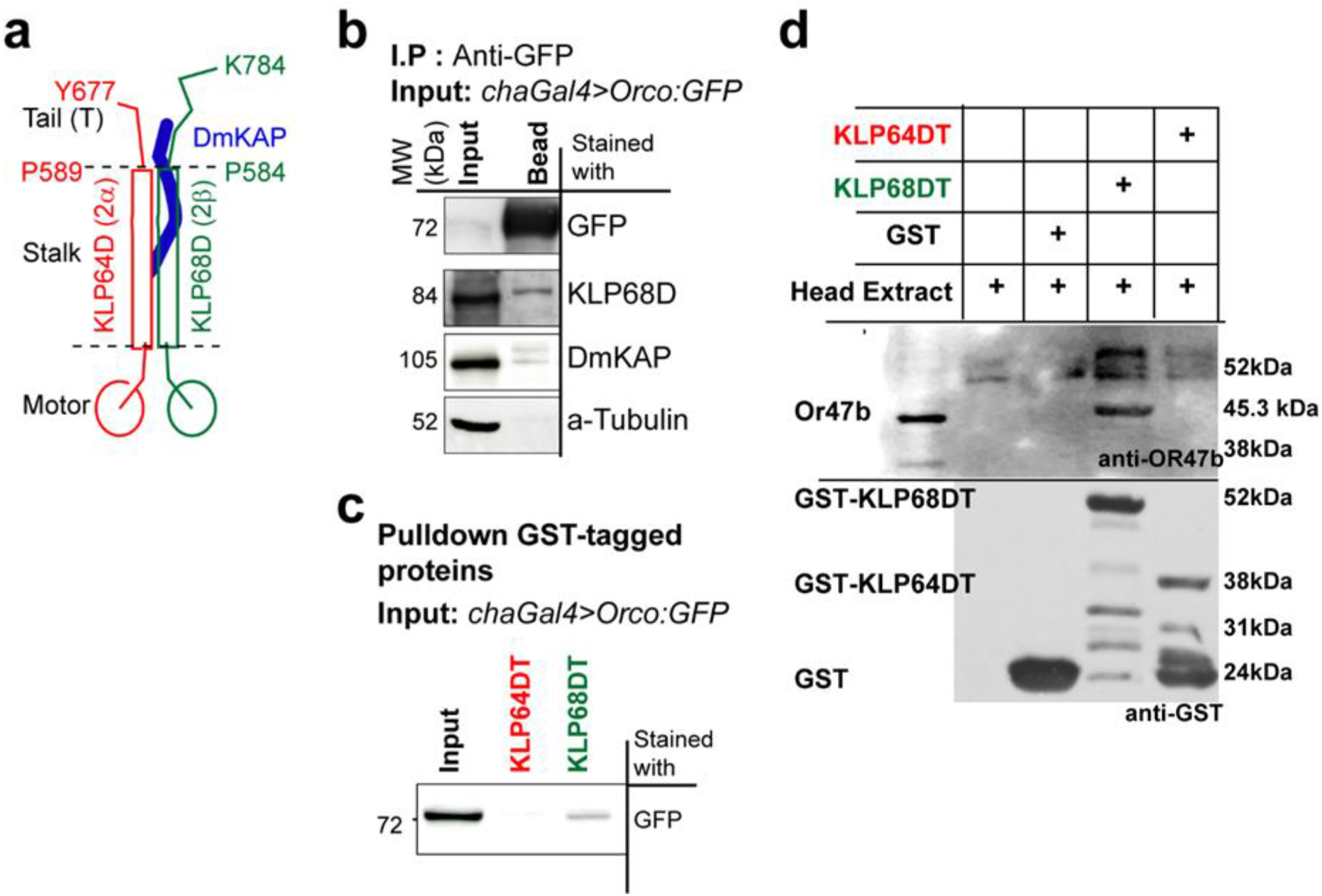
Orco associates with the C-terminal’ tail’ domain of the Klp68D (kinesin-2β) motor subunit. **a)** Schematic illustrates the organisation of *Drosophila* kinesin-2 and the tail domains of the motor subunits. **b)** Co-immunoprecipitation of the *Drosophila* kinesin-2 subunits from the head extracts of flies expressing Orco:GFP (driven by *chaGal4*) using an anti-GFP antibody. **c, d)** Affinity copurification of Orco::GFP (**c**), and OR47b (**d**), respectively, from the adult head extracts by using the GST tagged kinesin-2 tail fragments, KLP64D-T and KLP68D-T.

### Orco enriches in a characteristic distal domain at the ciliary OS

Next, we examined the Orco movement in the cilia after its entry. Due to the weak fluorescence of Endo-Ocro:GFP, extensive photobleaching and cuticular interference, the time-lapse imaging was not possible. The time-resolved plot profile analysis revealed that Orco is mostly enriched in the distal OS, in an approximately 4 μm band with a stereotypic peak at the distal OS (Figure 6a-b). This portion of the cilia, housed within the porous shaft of basiconica, consists of predominantly singlet microtubule-bearing branches (Shanbhag et al., 1999; Jana et al., 2011). Similar Orco localisation, distinct from that of the tubulin, was observed in the or42b cilium (Figure 6c, d). The distribution of Orco was maintained throughout 0-6 h AE, and it was distinct from that of the membrane marked by Bodipy-FL-C_12_ (Figure 6e, f). The cytoskeleton marked by *chaGal4>UAS-GFP:tubulin84B* and endogenous Jupiter:GFP (MAP) (Figure S6b), as well as the cytoplasm labelled by eGFP (Figure S6c), also elicited a relatively uniform distribution profile throughout the OS (Figure 6f, and S6). Thus, the enrichment domain appeared exclusive for Orco. Interestingly, the mCD8:GFP, a non-specific transmembrane protein, also found to have a distribution profile similar to that of Endo-Orco:GFP (Figure S6c), suggesting that transmembrane proteins are likely to be enriched in the distal OS domain of the olfactory cilia.

**Figure 6:**
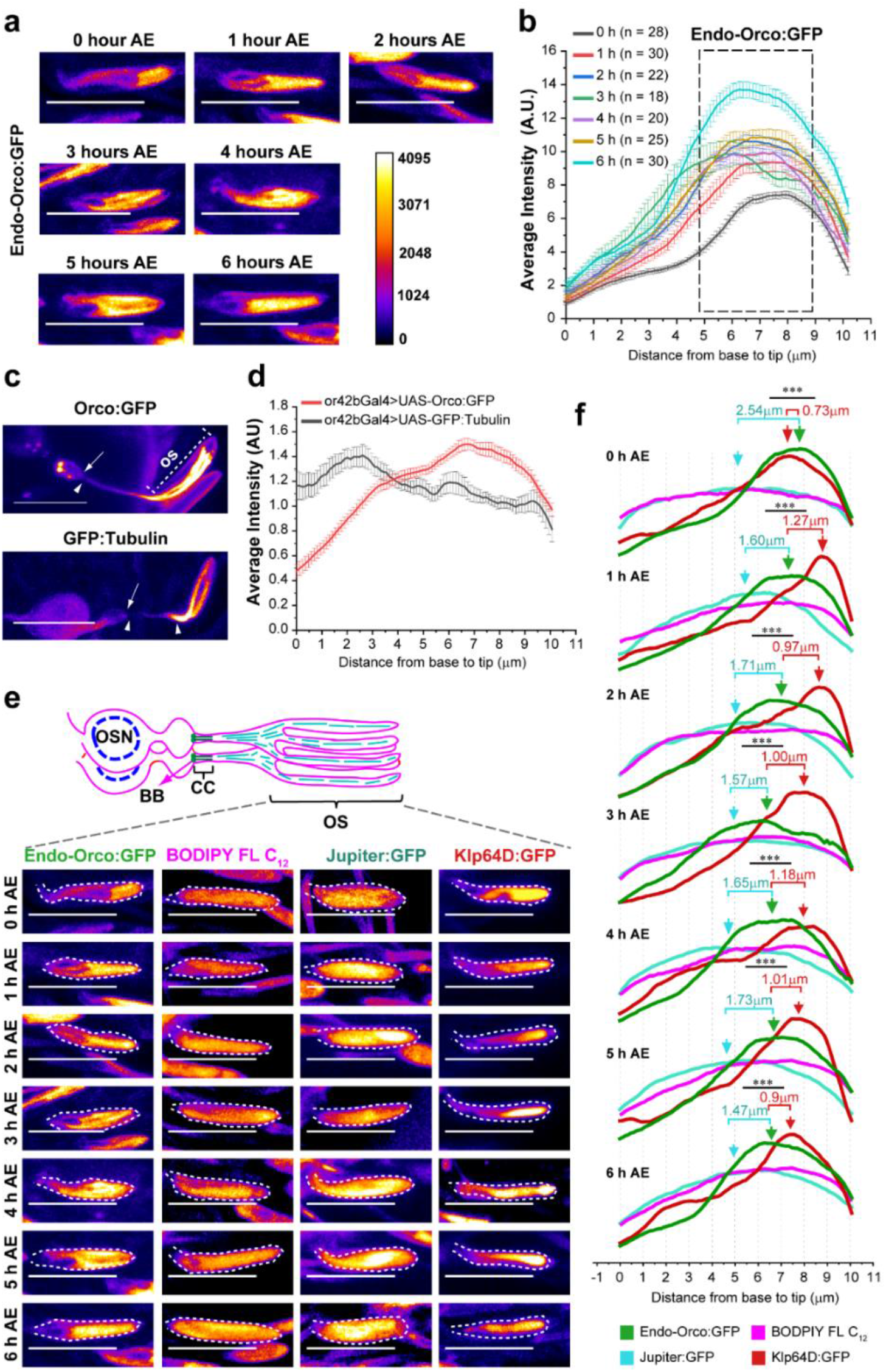
Orco enrichment domain in the cilia. **a, b)** Fluorescence micrographs (**a)** and plot profiles (**b)** of Endo-Orco:GFP distribution (average + S.E.M.) along the ciliary OS inside ab1-type *s. basiconica* during 0-6 hours AE. **c, d)** 5xGFP:Orco distribution along a single cilium expressed on or42b neuron inside ab1-type sensilla. **e, f)** Fluorescence micrographs **(e)** and relative distribution profiles **(f)** of Endo-Orco:GFP, Klp64D:GFP, Jupiter:GFP and BODIPY FL C_12_ along the ciliary OS inside ab1-type *s. basiconica* during 0-6 hours AE. Broken line marks the cuticle contour. The peaks of Endo-Orco:GFP, Klp64D:GFP and Jupiter:GFP are marked with green, red, and cerulean arrows, respectively. The separations between the peak Endo-Orco:GFP with Klp64D:GFP and Endo-Orco:GFP with Jupiter:GFP are indicated by red and cerulean numerals, respectively. The pairwise significance of difference was estimated using two-tailed Student’s T-test, p-values (*p < 0.05, **p < 0.01, and ***p<0.001) are shown on the plots.

Kinesin-2 motor interacts with Orco and facilitates the entry of Orco into the ciliary OS. Also, the kinesin-2 subunit Klp64D:GFP had a distribution profile that nearly overlapped with Endo-Orco:GFP at 6 hours AE (Figure S6d). Therefore, to comprehend if kinesin-2 aids the establishment of Orco domain in the ciliary OS, we estimated the separation between the profiles of Endo-Orco:GFP with that of Klp64D:GFP and Jupiter:GFP, respectively, at 0-6 hours AE. The Orco band partly coincided with that of the Kinesin-2 subunit Klp64D:GFP at different stages from 0-6 h AE (Figure 6f). Also, the Endo-Orco:GFP peak was much closer to that of the Klp64D:GFP as compared to the Jupiter:GFP. The analysis showed that kinesin-2 is selectively enriched at the OS domain along with the Orco. Therefore, kinesin-dependent transport could accumulate Orco in the distal part. To test the role of IFT in this process, we examined the localisation of the IFT-72 orthologue, Oseg-2:GFP (Avidor-Reiss et al., 2004), which was highly enriched in the IS (Figure S6a), and mostly uniform in the OS (Figure S8). We reason that the Orco enrichment would be executed independently of the cytoskeleton and bulk membrane transport.

### Kinesin-2 is required for the enrichment of Orco in the distal domain

To probe the possibility that kinesin-2 motor could enable the Orco enrichment in the distal OS domain, we estimated the total as well as relative intensity distribution of Endo-Orco:GFP along the cilia in control, *Klp64D^kj353^* mutant and kinesin-2 RNAi backgrounds during 0-6-h AE. The surface plots show that in the *Klp64D^kj353^* mutant, the levels of Endo-Orco:GFP is reduced (Figure 7a). In the kinesin-2 RNAi backgrounds, the Endo-Orco:GFP fluorescence was highly depleted from the OS domain, particularly at the 3-h and 6-h AE, intervening the active transport spurts at 1-2 h and 4-5 h AE, respectively (Figure 7a). Also, the full-width half maxima (FWHM) analysis of the Endo-Orco:GFP profile suggested a proximal spreading of the domain (Figure 7b). Note that dotted lines indicating the FWHM ingresses beyond the 4μm boundary. We also noticed that initially, during 0-3-h AE, the Endo-Orco:GFP distribution was proximally shifted by a significant margin in the *Klp64D^kj353^* background (Figure S6e), which was prominent in the kinesin-2 RNAi backgrounds at 6-h AE (Figure 7c), suggesting a comparatively higher penetrance of the Klp64D and Klp68D RNAi as the time progressed. Therefore, we concluded that interaction between Orco/ORx and kinesin-2 could maintain them in the designated domain at the ciliary OS.

**Figure 7:**
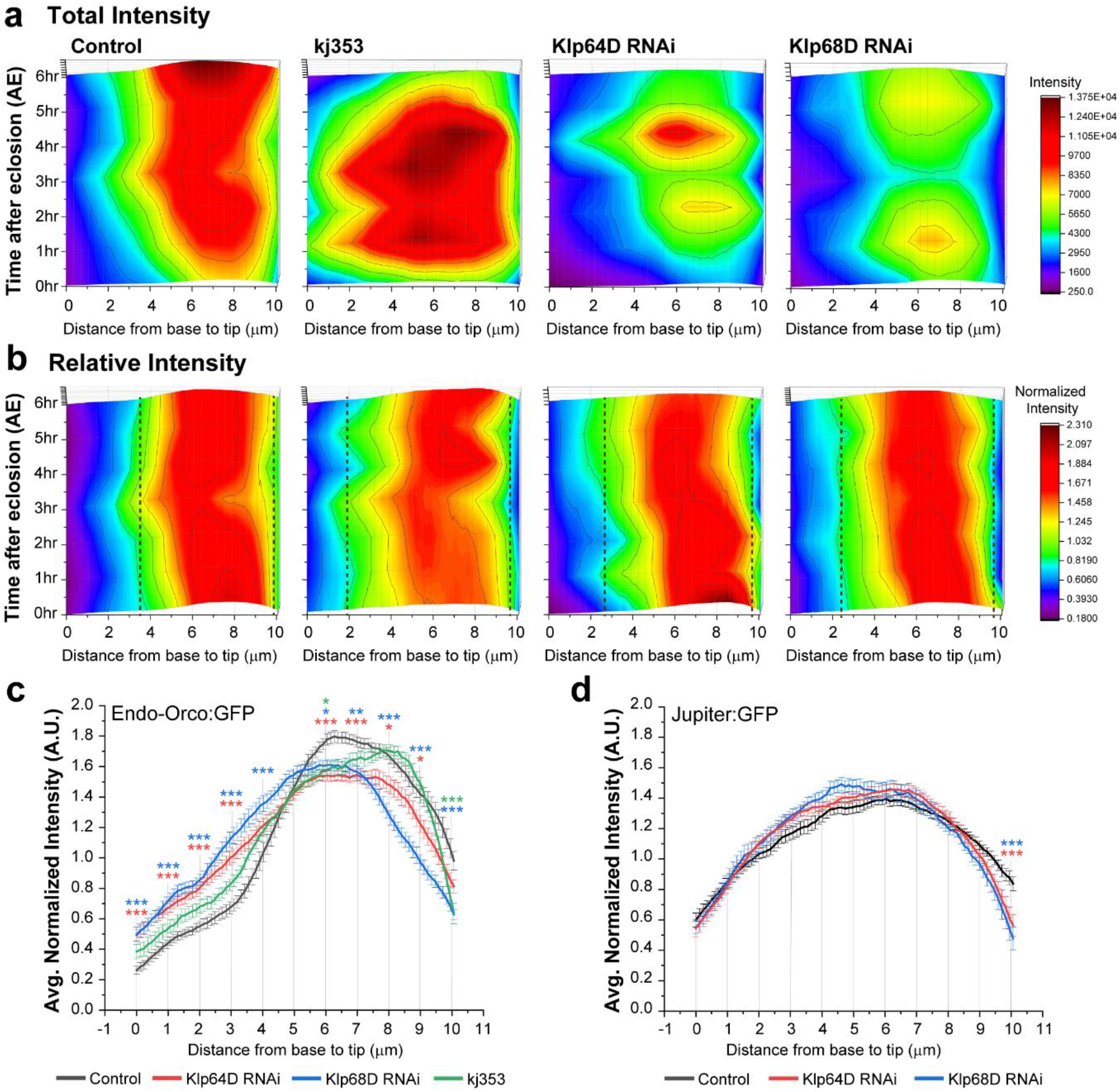
Endo-Orco:GFP distribution in kinesin-2 knockdown and mutant backgrounds. **a, b)** Surface plots for total **(a)** and normalised intensity **(b)** distributions of Endo-Orco:GFP along the ciliary OS during 0-6 hours AE in the wild-type control, homozygous *Klp64D^kj353^* mutant, and Klp64D and Klp68D RNAi backgrounds. Dotted lines demarcate the FWHMs of the Endo-Orco:GFP localisation domains. **c, d)** Comparisons of the Endo-Orco:GFP **(c)** and Jupiter:GFP **(d)** distribution (average + S.E.M.) along the ciliary OS at 6 hours AE. The pairwise significance of difference was estimated using one-way ANOVA test, p-values (*p < 0.05, **p < 0.01, and ***p<0.001) are shown on the plots.

## Discussion

The Orco/ORx localisation into the olfactory cilia is critical for the sensory reception (Benton et al., 2006) (Larsson et al., 2004). The cuticle shaft of the sensilla is porous, and it houses the ciliary outer-segment, which contains the branched extensions of cilia (Jana et al., 2011; Shanbhag et al., 2000). Here, we showed that the Orco is specifically enriched in the part of the branched OS that is housed inside the porous cuticle shaft. The branching increases the surface area of the cilia and accommodates a larger number of the receptors. In addition, packing a large number of branches, typically between 100-140 inside an LB-type shaft (~25-35 per cilium) would increase the receptor density and facilitate the trapping of cognate odorant molecules at very low concentrations. Hence, branching the outer segment and positioning Orco along the branches becomes critical for the optimum olfactory response. We found that the heterotrimeric kinesin-2 (kinesin-2α/β & KAP) motor associates with Orco/ORx complex, help to transfer the protein through the transition zone, and plays a crucial role in localising Orco to a specific subdomain in the ciliary outer-segment. Thus, kinesin-2 activity is independently implicated in efficient odour reception and olfactory behaviour.

The homodimeric kinesin-2 transports transmembrane protein into the cilia independent of IFT (Jenkins et al., 2006). The heterotrimeric kinesin-2 is implicated in transporting Rhodopsin (Bhowmick et al., 2009; Trivedi et al., 2012) and Transient Receptor Potential Vanilloid (TRPV) channels (Qin et al., 2005), via the IFT, although a recent study has contested the claims and suggested that kinesin-2 and IFT function is majorly restricted to the ciliary axoneme assembly (Jiang et al., 2015). Also, the behaviour analysis in homozygous Ceklp11 and Cekap1 mutant *C. elegans* suggested that the heterotrimeric kinesin-2 could maintain the chemosensory functioning of the AWB and AWC cilia (Evans et al., 2006; Mukhopadhyay et al., 2007) through IFT and BBS/Rab8 transports (Mukhopadhyay et al., 2008). Both the heterotrimeric and homodimeric kinesin-2 subunits are involved in IFT dependent and independent processes. Hence, it is difficult to distinguish between these two by genetic analysis involving total-loss-of-function mutants or RNAi knockdown. Consistent with this, we have also observed substantial sensory defects in the recessive, lethal mutants of Klp64D, which was associated with severe cilia growth defect (Jana et al., 2011). The isolation of homozygous viable Klp64D mutants with selective olfaction defects and mostly intact cilia ultrastructure helped to identify an independent role of kinesin-2 in the Orco/ORx transport in *Drosophila*.

How does the same kinesin-2 complex independently engage in both IFT-based maintenance and Oroc/ORX transport? We believe that it happens through a time-gated process. The basiconic cilia fully develop during the pupal stages (Jana et al., 2011), and a previous study showed that ciliary tubulin is highly stable with very low turnover as compared to that of the kinesin-2 motor in the adult stage (Girotra et al., 2017). Whereas, the bulk of the Orco is transported into the cilia at the adult stage immediately after eclosion. Hence, the adult-specific RNAi of kinesin-2 in OSNs had a selectively identifiable impact on Orco transport. It is still unclear whether the process may involve the IFT complex, although we found very little Oseg2/IFT72 punctae in the OS region.

The sensory signal-dependent maintenance of the olfactory cilia morphology and chemosensory behaviour in *C. elegans* involves components of the Intraflagellar Transport (IFT), Bardet-Biedel Syndrome (BBS) proteins and Rab8 (Mukhopadhyay et al., 2008). The exact correlation between the receptor transport, cilia remodelling and sensory modulation is still unclear. We were intrigued to find that the Orco/ORx entry is specific to the cilia type, and it is tightly regulated during development. What might be the mechanism of such selectivity? Recent studies suggested that post-translational modifications such as SUMOylation trigger the receptor entry into the cilia (McIntyre et al., 2015). Also, concerted activation of tubby-like protein, exocyst complex and small GTPases like Rab8 regulate the delivery of transmembrane protein to the ciliary pocket at the transition zone. The Bardet-Biedel Syndrome (BBSome) proteins and the Rab-family of small GTPases plays a critical role in the second stage of this transport (Kuhns et al., 2019; Nachury et al., 2010). The entry and exit through the transition zone are also regulated by cell signalling (Corbit et al., 2005; Pal et al., 2016). Therefore, a combination of these functions could deliver Orco/ORx at the base of the cilia. The ciliary membrane is isolated from that of the plasma membrane through an elaborate structure, called ciliary sheet and ciliary necklace (Vieira et al., 2006), at the base of the cilia. Therefore, after the Orco/ORx carrying vesicles fuse at the ciliary pocket, these proteins need to be actively taken across the barrier. We showed that interaction with the heterotrimeric kinesin-2 motor would be responsible for this movement.

## Materials and Methods

### Drosophila culture and stocks

All the fly stocks used in this study are listed in Table supplement 1. Flies were reared on standard cornmeal agar medium at temperature 25 ^μ^C for all the studies. All the experiments were done on dissected antennae of adult flies of different ages as per the experimental requirements.

### Drosophila sample preparation

For live imaging of adult Drosophila antennae, antennae were dissected and mounted in a drop of grade 100 Halocarbon oil (Sigma^®^). For imaging of aged pupal antennae, late third instar larvae were monitored every 30 minutes until they become stationary and this was considered the beginning of the white pupae stage, marked as 0 hr APF, and maintained at 25°C afterwards. The pupal case was opened, and the head was dissected in Phosphate Buffered Saline (PBS: 137 mM NaCl, 2.7 mM KCl, 10 mM Na2HPO4, 1.76 mM KH2PO4 and pH 7.4) at designated times after the 0h APF. Once isolated, the heads were fixed in 4% formaldehyde solution (4% Paraformaldehyde in PBS, pH 7.4) for 30 minutes at room temperature, followed by five washes with PBS. The antennae bearing the second and third segments were then pinched off from the head and mounted in a drop of Vectashield^®^ (Vector Laboratories Inc., USA) on a glass slide under a 0.17 mm coverslip.

### Immunostaining

For Orco immunostaining, 1 day old adult Drosophila heads were dissected, sectioned in a cryo-microtome (Leica, GmBh) and fixed following standardised protocols. 10 μm sections were laid on a poly-D-Lysine coated slide 1 and fixed in 4% formaldehyde solution for 30 minutes at room temperature. Then, they were stained using 1:50 dilutions of MAb-Orco (Abcam, USA) and Alexa647-conjugated goat anti-mouse (Molecular Probes Inc., USA) following the published method (Dobritsa et al., 2003).

### Dyes microinjection

Both Alexa 546-Dextran (Invitrogen, USA) and Bodipy FL C_12_ (Invitrogen, USA) dye were injected into the head capsule of anesthetised adults and incubated for 10 minutes to allow the dye to permeate the third antennal segment through hemolymph. Exclusion of Alexa 546-Dextran from the sensilla confirmed that the ciliary lymph is isolated from the hemolymph (Figure S7). Therefore, we concluded that the ciliary membrane exclusively contributes to the BOPIPY label in the sensillum shaft (Figure S6c).

### Image Acquisition and analysis

All fluorescence images were obtained under constant acquisition conditions in Olympus (Olympus Imaging Corp., Japan) confocal microscope FV1000SPD using a 63x oil 1.4 NA objective lens or FV3000 using a 60x oil 1.4 NA objective lens. Subsequently, all images were processed using ImageJ^®^ (rsweb.nih.gov/1 IJ). Detailed analysis for volume and intensity measurements on 3-D volume-rendered image stacks was done using Imaris^®^ 6.1.5 graphics package (Bitplane Scientific Solutions AG, Switzerland). All figures containing images and line arts were composed in Illustrator CS6^®^ (Adobe Inc., USA). The cilia of the ORNs innervating a single sensillum are projected together into the sensory shafts. They are tightly intertwined and could not be resolved using confocal microscopy. Therefore, all the length and volume measurements indicate the combined value of the cilia inside a shaft.

### Electro antennogram recording

As described before (Jana et al., 2011).

### Electron Microscopy

As described before (Jana et al., 2011).

### Drosophila head extracts

Fly heads were manually separated by razor blades, and homogenised in 250 μl of ice-cold, high KCl buffer [19 mM KH2PO4, 81 mM K2HPO4, 400 mM KCl and 1 mM DTT (pH 7.4)], containing the protease inhibitor cocktail (Roche GmbH). Motorised plastic homogeniser and specially designed microfuge tubes were used for this purpose (Pellet PestleR, Kontes, USA). The homogenate was first centrifuged at 17,000 x g for 30 minutes in a refrigerated micro-centrifuge (Biofuge Fresco, Heraeus AG, Germany) to clear the tissue debris and then the supernatant was further centrifuged at 100,000 x g for 1 hour at 4⁰C to separate the soluble (supernatant) and membrane-associated proteins (pellet). To extract the membrane-associated proteins, the pellet was homogenised 1 in 250 μl high KCl buffer containing 1% Triton-X-100 and 1% Sarkosyl, then centrifuged at 100,000 x g. The resultant supernatant was used as the detergent extracted fraction of the membrane pellet for subsequent pulldown experiments.

### Affinity copurification

100 μl aliquots of the soluble and the detergent extracted fractions of the Drosophila head homogenates were separately mixed with fixed amounts of purified recombinant proteins in the high KCl buffer. The mixtures containing GST-tagged recombinant proteins were incubated with an equal quantity of Glutathione SepharoseTM 8 beads for 2 hours at 4°C. Following this, the Glutathione SepharoseTM beads were washed with several changes of the ice-cold, high KCl buffer containing 0.1 mM Glutathione. Finally, the bound proteins were eluted by incubating the beads in 100 μl proportions of the high KCl buffer containing 1 mM Glutathione. 0.3 ml of the eluted fractions was further incubated with 0.3ml Glutathione SepharoseTM beads and washed and eluted with equal bed volumes of TMN-D containing 1 mM Glutathione. The buffer conditions and salt concentrations were maintained constant throughout this process. The beads were boiled in sample loading buffer and loaded on SDS-PAGE and transferred to PVDF membrane.

### Western blots and immunostaining

The protein mixtures were separated on SDS-PAGE and transferred to PVDF membrane (Hybond-P, Amersham Biosciences Plc. UK) following the supplier’s protocol and incubated in different primary antisera solutions (1:1000 dilutions) in 20 mM Tris-buffered saline (TBS, pH 7.4) containing 0.1% TweenR 20. Subsequently, they were incubated either in Rabbit anti-mouse:HRP (dilution, 1:20000; Sigma Chemicals Co. MO, USA), or Goat anti-rabbit:HRP (dilution, 1:20000; Sigma Chemicals Co, MO, USA), in the TBS-T, and developed by using ECLR chemiluminescence detection kit (GE Healthcare Ltd. USA).

### Statistical Analysisxs

All statistical comparisons were carried out using either student’s t-test or one way ANOVA with p values calculated according to Bonferroni Test in Origin^®^. P values for data sets in each figure are indicated in the respective figure legends.

## Acknowledgement

This manuscript is dedicated to the memory of the late Prof. Obaid Siddiqui. We thank Prof. Tomer Avidor-Reiss, University of Toledo, Toledo, USA, for the UAS-Oseg-2:GFP fly stock, B. Karmakar, and R. Phadke for technical help with the instrumentation and experiments, and L. Borde for assistance with the TEM. We especially acknowledge M. Bettencourt-Dias, IGC, Portugal, for exceptionally generous support. A part of the experiments was carried out at IGC by SCJ.

## Author’s contribution

**SCJ, AS, AJ, MG** - isolation and phenotypic characterisation of kinesin-2 mutants; **PD, AJ** - Orco transport analysis; **PD** - analysis of Orco subdomain enrichment and mechanism; **SCJ, MG** - copurification assays; **SCJ, SS** - TEM data; **KR, SCJ, AJ, PD** – experimental design, data analysis and manuscript writing. **SCJ** and **KR** evolved the concept.

