## Supplemental Figures for "Heterotrimeric kinesin-2 motor subunit, KLP68D, localises *Drosophila* odour receptor coreceptor in the distal domain of the olfactory cilia"

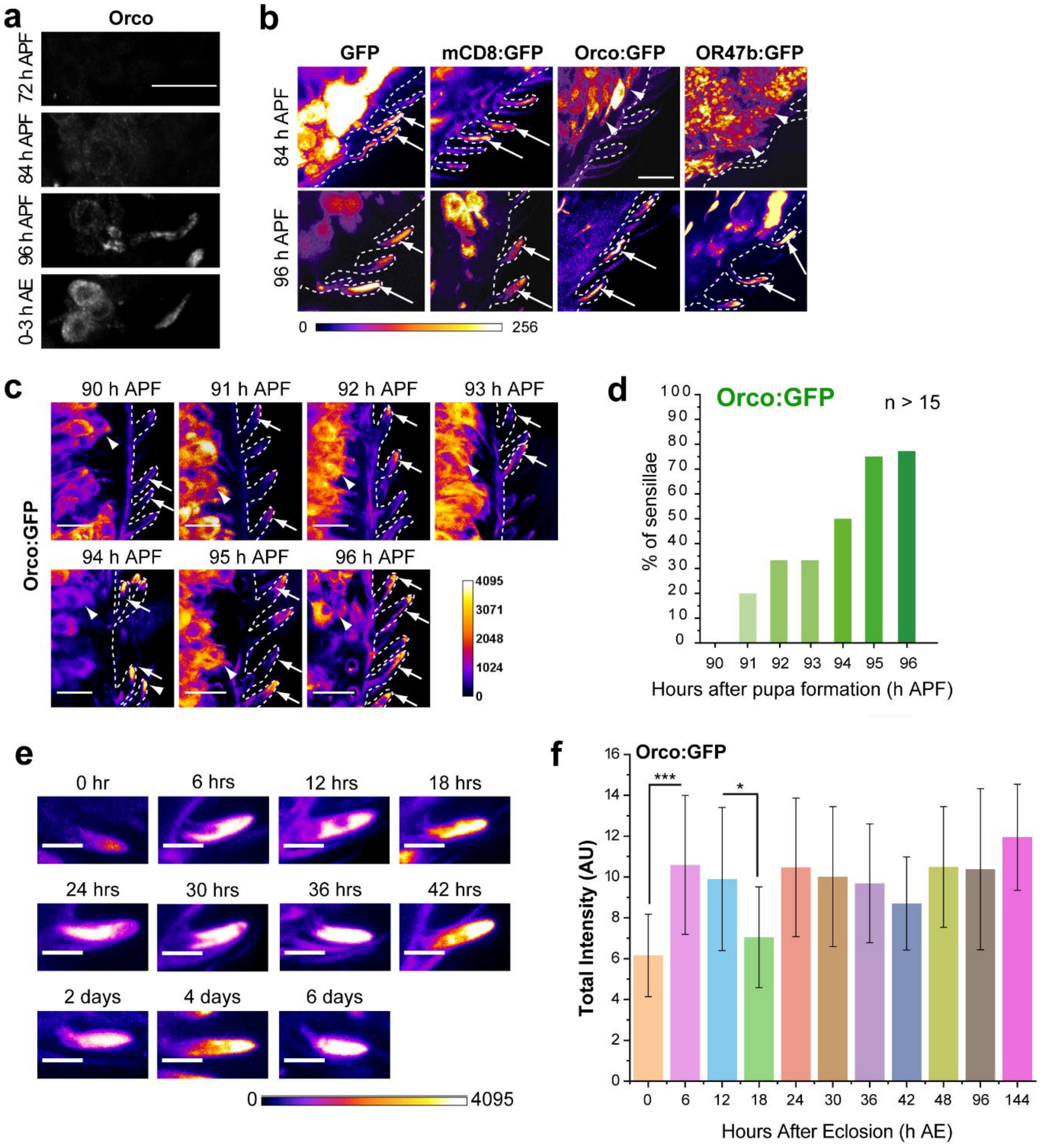


**<u>Figure S1 (linked to Figure 1):</u> Developmental profile of OR localization in the cilia on OSNs innervating s. basiconica. a)** Immunostaining using anti-Orco antibody showing Orco localization in cilia in s. basiconica of pupae during 72 - 96 h APF and 0-3 h AE. b) GFP, mCD8:GFP, Orco:GFP and Or47b:GFP localization in cilia in s. basiconica during 84 - 96 h APF. Expression of all driven by *chaGal4*. **c)** Endo-Orco:GFP localization in pupal cilia in s. basiconica from 90 hour to 96 hours APF. Arrowheads indicate cell bodies of the OSNs and arrows mark the ciliary OS. **d)** Percentage of sensillae with Endo-Orco:GFP localization in the ciliary OS of pupal cilia in s. basiconica from 90 - 96 h APF. **e)** Endo-Orco:GFP localization in the ciliary OS in s. basiconica of adult flies during 0 h - 6 days AE. **f)** Total fluorescence intensity of Endo-Orco:GFP in the ciliary OS in s. basiconica of adult flies from 0 h to 6 days AE. The pairwise significance of difference was estimated using one-way ANOVA test, p-values (*p < 0.05, **p < 0.01, and ***p<0.001) are indicated on the plots. Error bars represent as <u>+</u> SD. All images are shown in the false colour intensity heat map (FIRE, ImageJ®). Scale for images **a), b)** and **c)** 10 µm; **e)** 5 µm.

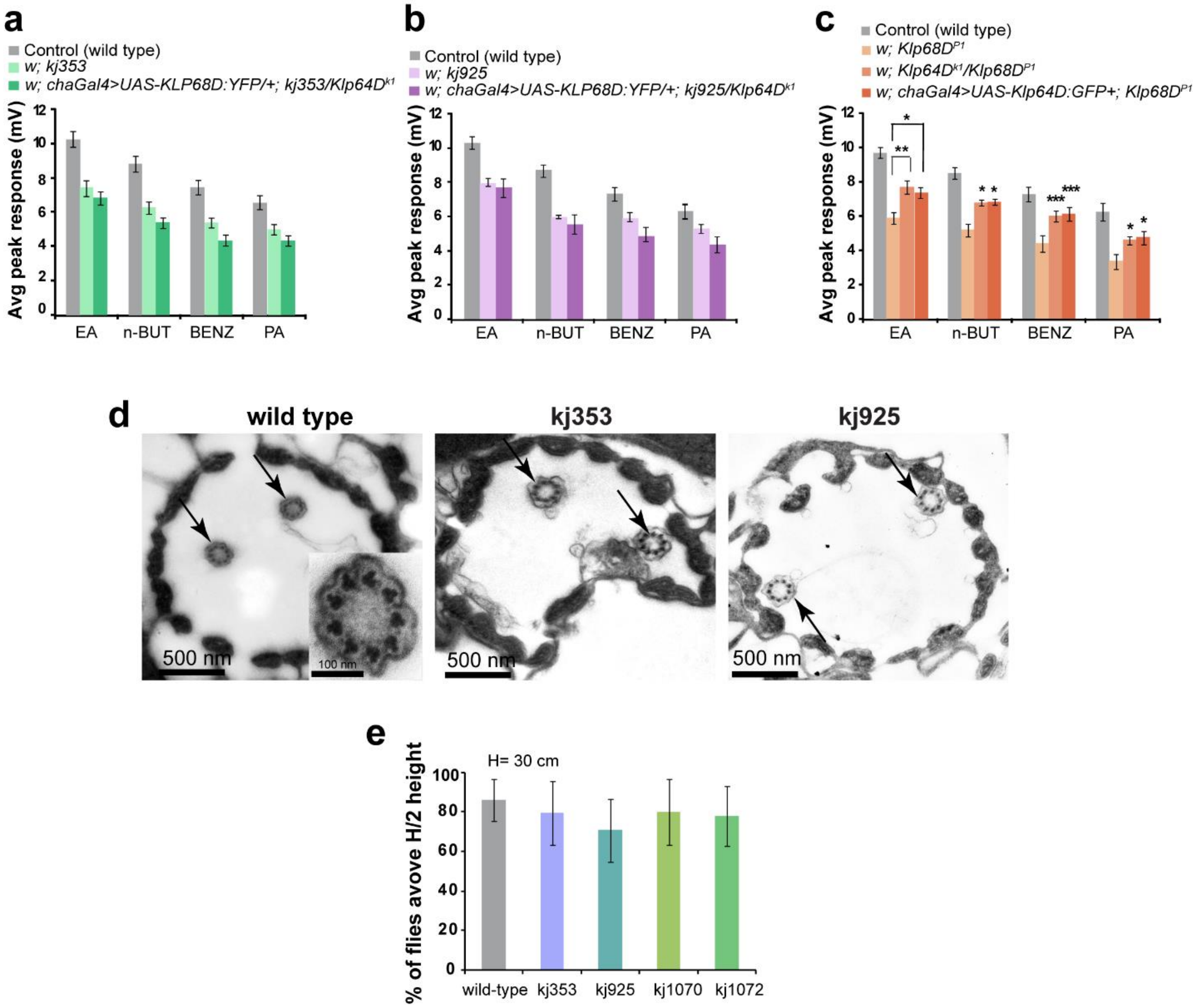


**Figure S2 (linked to Figure 2): *kj* alleles map to the Klp64D and not in the Klp68D locus. a-c) EAG response defects of the homozygous *Klp64D$^{kj}$* mutants are not rescued by the OSN-specific expression of the *KLP68D* transgene.** Histograms indicate average (+ SD, N ≥ 30) electroantennogram (EAG) responses from the antennae of various mutant combinations. *Klp64D$^{k1}$* is a lethal total-loss-function allele of *Klp64D* and *Klp68D$^{P1}$* is a hypomorphic, viable, P-element insertion allele of *Klp68D*. *chaGal4* induced expression of *UAS-KLP68D-YFP*, as well as *UAS-KLP64D-GFP* in OSNs rescued the cilia development and EAG defects of the homozygous *Klp68D$^{P1}$* (Jana et al., 2011). The data shown in panel c indicate that loss of KLP68D could partly compensate by overexpression of KLP64D. **d-e) The chordotonal cilia are unaffected in homozygous Klp64D$^{kj}$ backgrounds. d)** TEM images of cross sections of scholopedia from Johnston's Organs in wild-type and homozygous mutant antennae. Each scholopedium contained two sensory cilia (arrows), with 9+0 organization of microtubule (inset). The chordotonal cilia appeared normal in the mutants. The chordotonal cilia play important roles in proprioception and anti-genotropic walk. **e) Homozygous *Klp64D$^{kj}$* alleles are coordinated.** Negative geotaxis (test of coordination) of wild-type and homozygous *kj* adults was measured by estimating the relative number of flies above the half-length of a 30 cm cylinder 5 minutes after banging. Homozygous *w1118* was used as the wild-type control. It showed that they were coordinated like wild type.

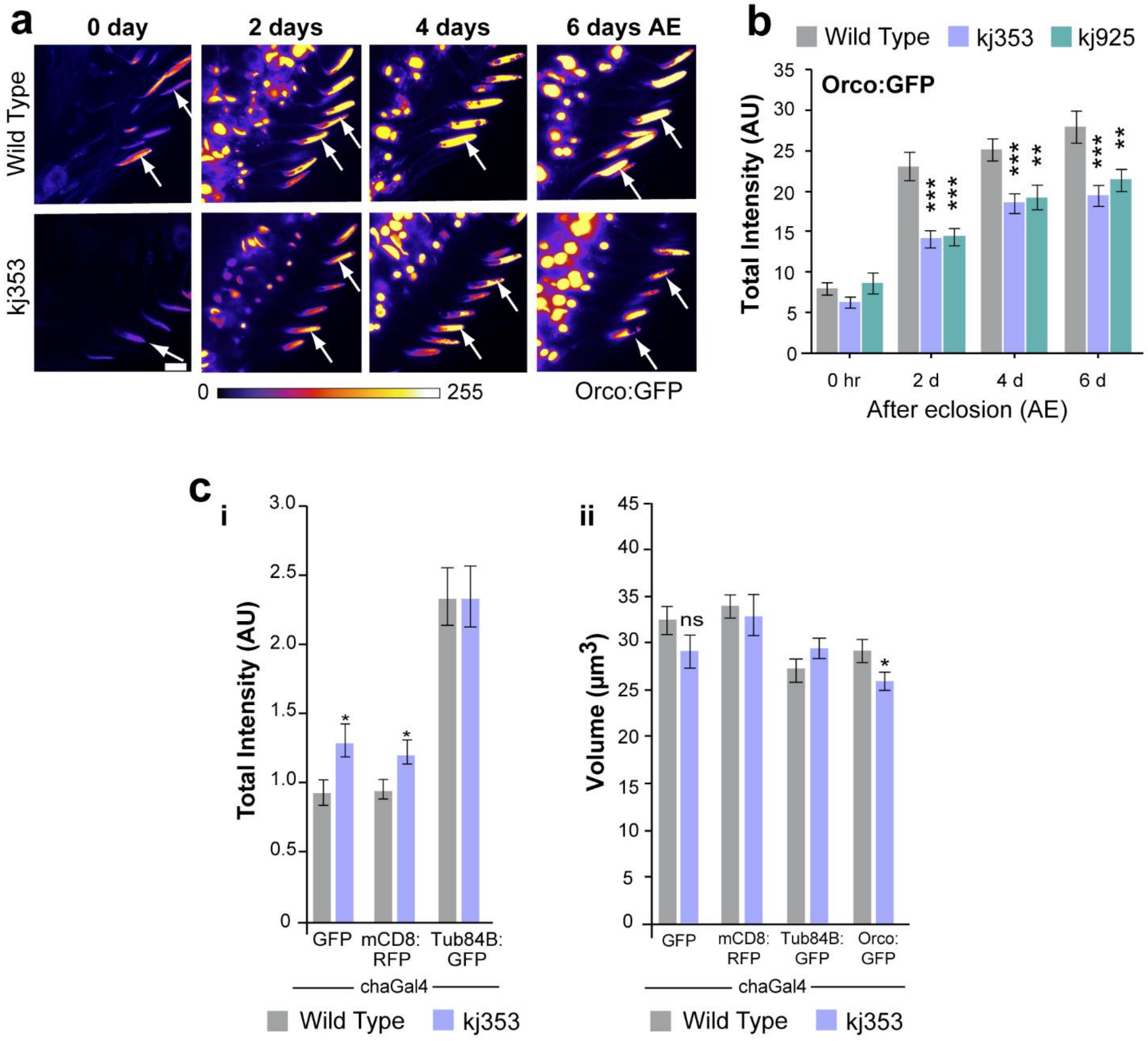


**Figure S3 (linked to Figure 3): Orco:GFP localization in the ciliary OS and ciliary structure characterization of Klp64D kj353 and kj925 mutants. a)** Reduced localization of Orco:GFP (expressed by *chaGal4*) in the ciliary OS in s. basiconica of wild type and Klp64D kj353 and kj925 mutants from 0 - 6 days AE. b) Total fluorescence intensity of Orco:GFP (driven by *orcoGal4*) in the ciliary OS in s. basiconica of wild type and Klp64D kj353 and kj925 mutants. The pairwise significance of difference was estimated using two-tailed Student's T-test, p-values (*p < 0.05, **p < 0.01, and ***p<0.001) are indicated on the bars. Error bars represent as + SEM. All images are shown in the false colour intensity heat map (FIRE, ImageJ®). Scale for images- 10 µm. **c-i, ii)** Total fluorescence intensity and ciliary volume of ciliary OS in s. basiconica marked by soluble protein- EGFP, membrane- mCD8:GFP, cytoskeleton- Tubulin::GFP and Orco:GFP (all driven by *chaGal4*) of two day AE control and Klp64D kj353 mutant flies. The pairwise significance of difference was estimated using two-tailed Student's T-test, p-values (*p < 0.05, **p < 0.01, and ***p<0.001) are indicated on the bars. Error bars represent as + SEM. d)

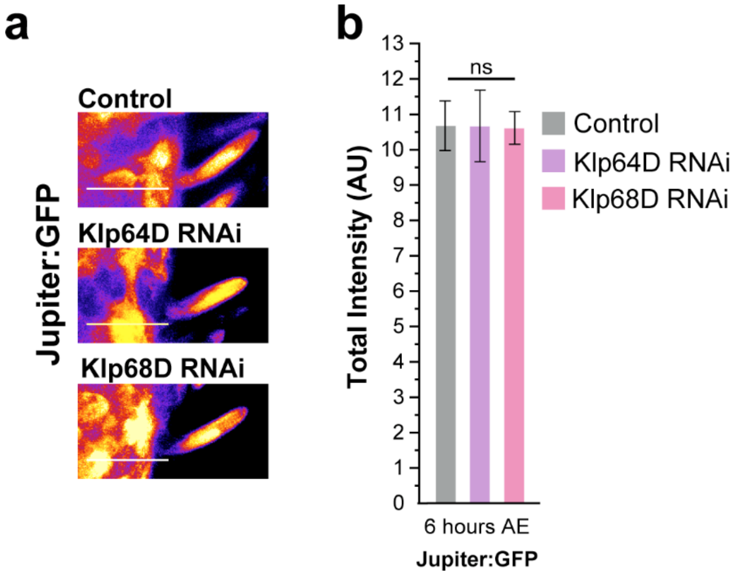


<u>Figure S4 (linked to Figure 4):</u> **Cytoskeleton of the cilia in s. basiconica is unaffected in upon adult specific knock down of kinesin-2 subunits, Klp64D and Klp68D. a)** Jupiter:GFP localization in the ab1-type s.basiconica from control (*or83bGal4/+; Jupiter:GFP/UAS-Dicer*), Klp64D RNAi (*or83bGal4/UAS-Klp64D RNAi; Jupiter:GFP/UAS-Dicer*) and Klp68D RNAi (*or83bGal4/UAS-Klp68D RNAi; Jupiter:GFP/Uas-Dicer*) background at 0 hour and 6 hours AE. **b)** Total fluorescence intensity of Jupiter:GFP in Klp64D RNAi and Klp68D RNAi background at 0 hour and 6 hours AE. The pairwise significance of difference was estimated using one-way ANOVA test. Error bars represent as $\pm$ S.D. All images are shown in false colour intensity heat map (FIRE, ImageJ®). Scale for images - 10 µm.

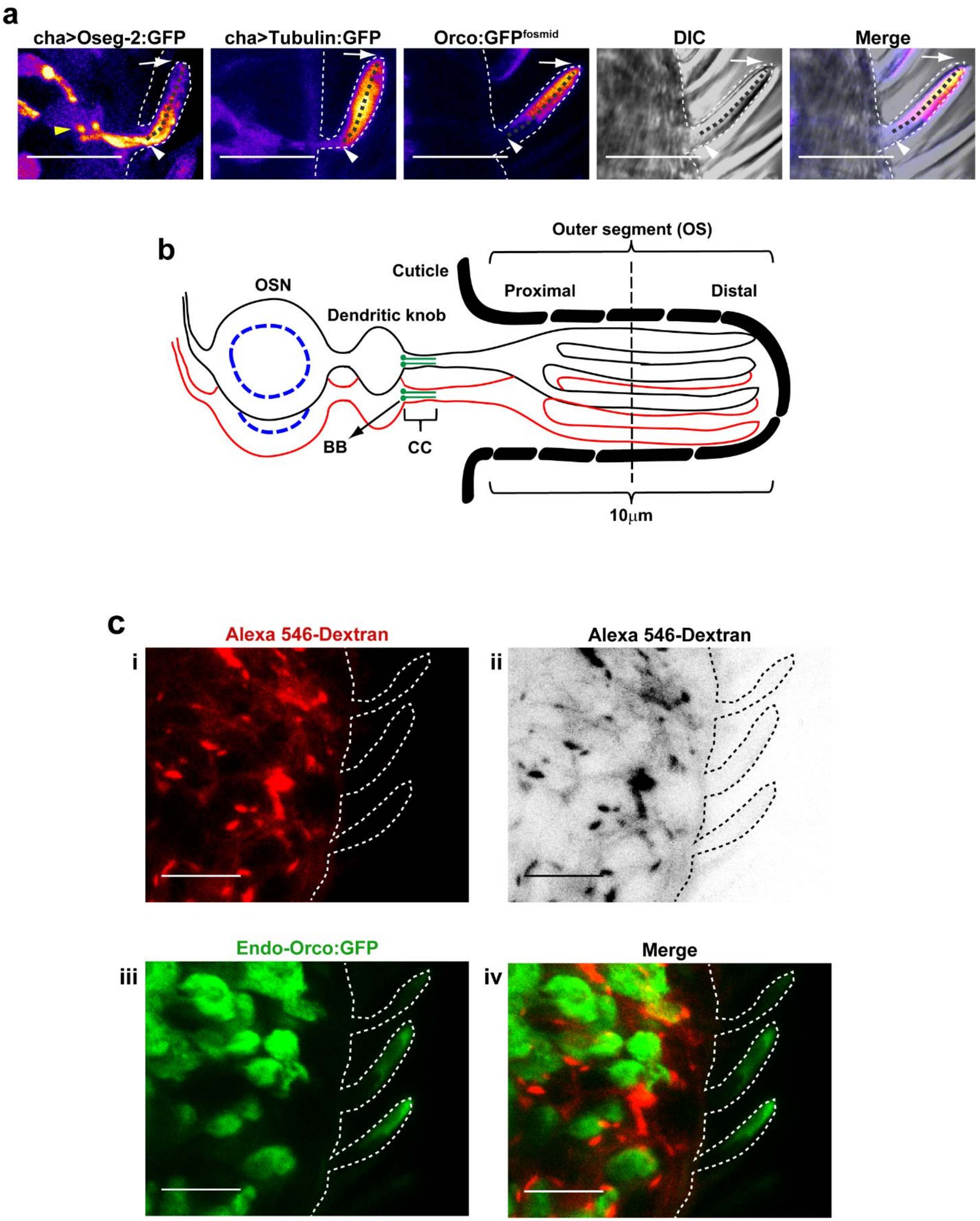


<u>**Figure S5 (linked to Figure 6):**</u> **Estimating the length of the cilia to measure the plot profiles**. **a)** Fluorescence micrographs of cilia in *s. bascionica* showing localization of Oseg-2:GFP, Tubulin:GFP, Endo-Orco:GFP, DIC of the cuticle of *s. basiconica*, and merge of DIC and Endo-Orco:GFP. The starting point of the ciliary OS (white arrowheads) was decided by the Oseg-2:GFP distribution and start point of the Endo-Orco:GFP. Feeble amount of Endo-Orco:GFP localize at the inner segment. It majorly starts to localize from the end of the inner segment and the start of the outer segment (white arrowhead). Oseg-2:GFP is found to localize from the basal body (yellow arrowhead) followed by the inner segment till major part of the middle region of the ciliary OS, however, sparsely localizing the distal end. The distal

most end of the ciliary OS marks the endpoint (white arrows) for measurement of the plot profiles. The length along which the line segment was drawn to measure the plot profiles is represented by a dark grey dotted line. All images are shown in the false colour intensity heat map (FIRE, ImageJ®). Scale for images – 10 µm. **b)** Schematic of an *s. basiconica* depicting the ciliary structure and the domain of the ciliary OS which measures to be approximately 10 µm. **c) Localization of Alexa 546-Dextran is excluded from the OSNs and sensillae. i)** Alexa 546-Dextran marking the free space in the third antennal segment. **ii)** Inverted image of Alexa 546-Dextran. **iii)** Endo-Orco:GFP marking the OSNs and the s. basiconica. **iv)** Merge of Alexa 546-Dextran and Endo-Orco:GFP depicting that Dextran is excluded from the lymph surrounding the olfactory sensory cilia innervating the sensillae. Scale for images - 10 µm.

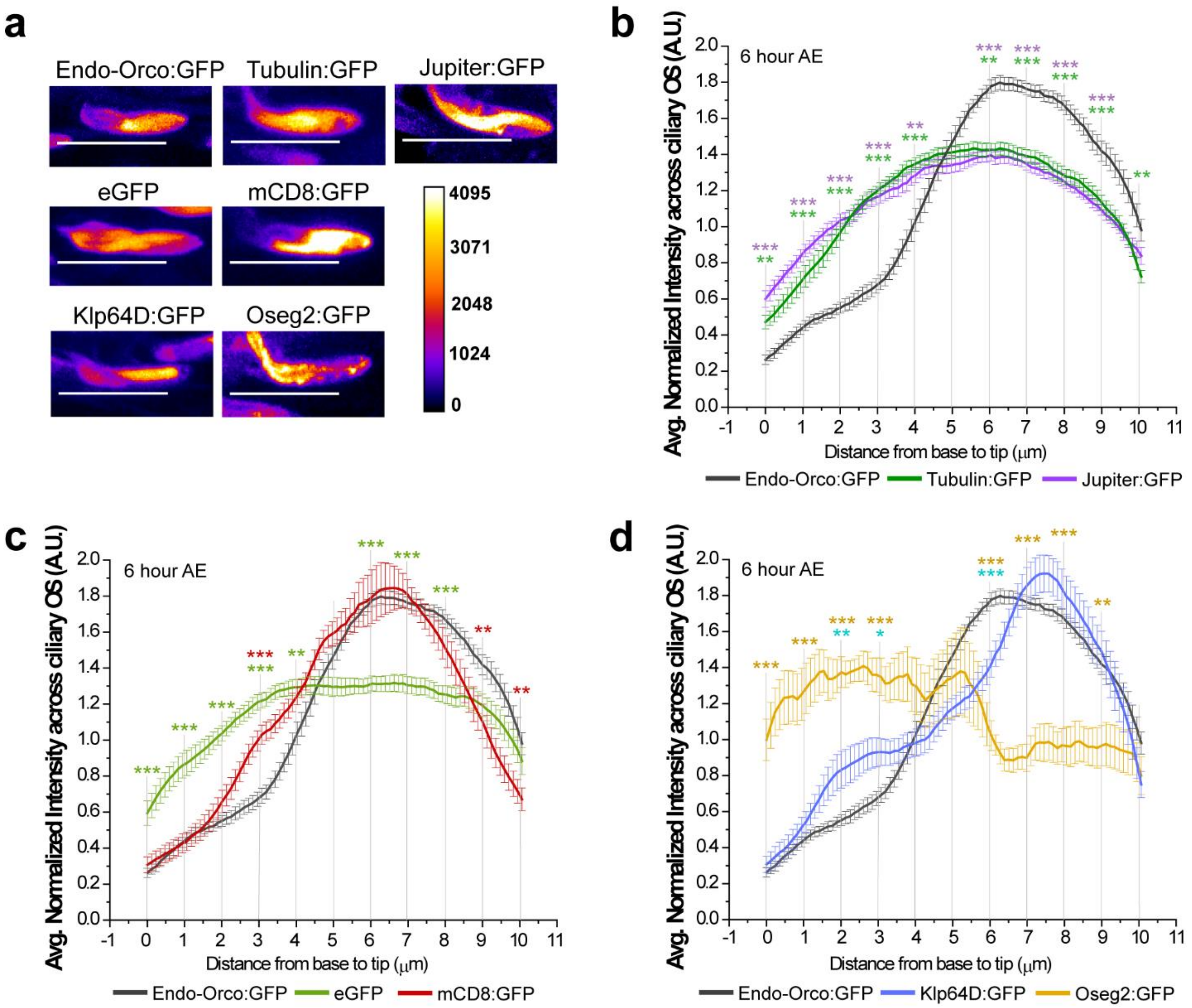


**<u>Figure S6 (linked to Figure 6):</u> Comparison of the domain of Orco accumulation with that of cytoskeleton, membrane and kinesin-2 motor. a)** Ciliary localization of Endo-Orco:GFP, cytoskeleton markers-Tubulin:GFP and Jupiter:GFP; soluble protein- EGFP; membrane marker- mCD8:GFP; and ciliary trafficking machinery- Klp64D:GFP and Oseg2:GFP, at 6 h AE. Except for Jupiter:GFP (protein trap) and Endo-Orco:GFP (fosmid line), the transgene expressions were driven by the *chaGal4*. **b-d)** Plot profiles of the distribution (average ± SEM) of Endo-Orco:GFP and the above ciliary markers across the ciliary OS in ab1-type s. basiconica of 6 h AE. The pairwise significance of difference was estimated at 1 [illegible]m gap along the length of the cilia using one-way ANOVA test, p-values (*p < 0.05, **p < 0.01, and ***p<0.001) are indicated on the plots. All images are shown in the false colour intensity heat map (FIRE, ImageJ®). Scale for images- 10 µm.

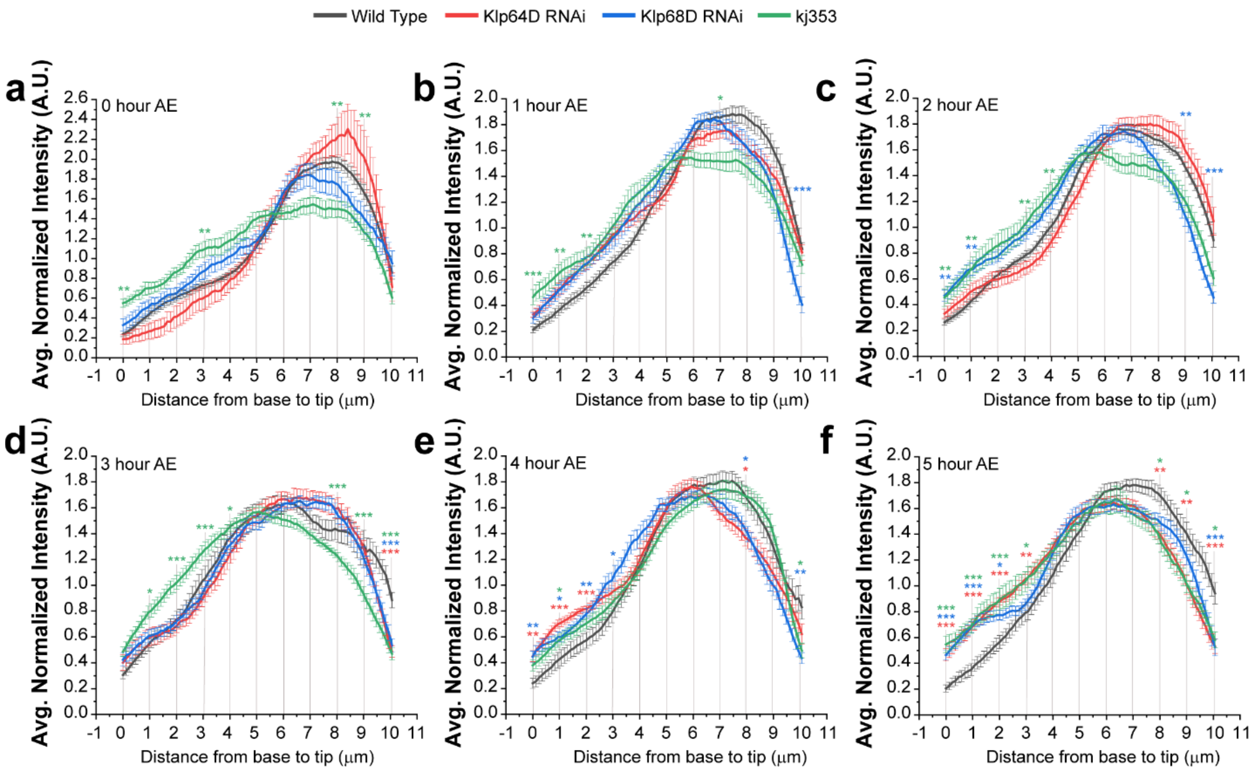


**Figure S7 (linked to Figure 7): Loss of kinesin-2 shifts the Endo-Orco:GFP enrichment towards the proximal domain. a)** Fluorescence intensity heat map of Endo-Orco:GFP along the ciliary OS of ab1-type *s. basiconica* during 0-5 h AE due to the knock-down of kinesin-2 motor subunits – KLP64D and KLP68D. and in the homozygous $Klp64D^{kj353}$ backgrounds, respectively (scale ~10 µm). **b-g)** Comparison of the relative distribution of Endo-Orco:GFP during 0 -5 h AE in control, KLP64D RNAi, KLP68D RNAi, and homozygous Klp64D kj353 mutant backgrounds. The pairwise significance of the differences was estimated at a few sample points along the ciliary OS using one-way ANOVA test, p-values (*p < 0.05, **p < 0.01, and ***p<0.001) are indicated on the plots. Error bars represent as <u>+</u> SEM.
