## Supplemental Tables for "Heterotrimeric kinesin-2 motor subunit, KLP68D, localises *Drosophila* odour receptor coreceptor in the distal domain of the olfactory cilia"

**Table S1 (linked to Figure 2): Allelic complementation analysis of *kj* isolates with known *Klp64D* alleles and the transgenic rescue of the defect, estimated using elactroantennogram (EAG).**

| | | *Klp64D (recessive, uncoordinated alleles}* | | | | | | Control | |
|---|---|---|---|---|---|---|---|---|---|
| | | *k5* | | *l3* | | *l4* | | + | |
| | | EA | nBut | EA | nBut | EA | nBut | EA | nBut |
| *Klp64D (jump response defect, viable alleles )* | ***kj353*** | 6.7±0.5** | 4.2±0.2** | 6.2±0.3** | 3.9±0.5** | 7.5±0.8** | 5.4±0.6* | 9.0±0.6 | 7.8±0.9 |
| | ***kj429*** | 6.1±0.4** | 4.3±0.4** | 6.2±0.3** | 4.2±0.5** | 5.0±1.5* | 4.1±1.2* | 8.3±0.6 | 6.4±0.4 |
| | ***kj925*** | 4.5±1.2** | 3.0±0.6** | 6.0±0.3** | 3.6±0.4** | 6.2±0.8** | 3.8±0.7** | 9.5±1.3 | 7.2±1.2 |
| | ***kj1070*** | 5.7±0.6** | 3.0±0.8** | 5.6±0.4** | 3.7±0.4** | 6.3±1.4** | 3.5±0.5** | 8.5±0.9 | 6.5±0.6 |
| | ***kj1072*** | 5.9±0.8** | 3.6±0.8** | 5.5±0.3** | 3.8±0.3** | 6.3±0.7** | 3.5±0.3** | 9.7±0.6 | 7.6±0.9 |
| | **Rescue Genotypes** | | | | | | | EA | n-But |
| *Transgenic Rescue* | ***w; Gal4$^{OR83b}$/UAS-KLP64D; kj353*** | | | | | | | 8.7±0.9 | 7.9+0.3 |
| | ***w; Gal4$^{OR83b}$/UAS-KLP64D; kj925*** | | | | | | | 10.2+1.3 | 9.3+1.3 |
| | ***w; Gal4$^{SG18.1}$/UAS-KLP64D; kj353*** | | | | | | | 9.5+0.7 | 8.1+1.3 |
| | ***w; Gal4$^{SG18.1}$/UAS-KLP64D; kj925*** | | | | | | | 10.4+0.3 | 9.1+0.8 |

**Color key: Ethyl Acetate (EA), n-Butanol (n-But)**

*Note:* EAG values were obtained from 2-days old adult flies of specific genotypes. V*alues in each cell indicate average (± S.D.) responses in mV (N≥6) for each genotype. Pairwise significance of differences was calculated with respect to the wild type control values for the respective odours using two tailed Student's T-test. Significant (*, p<0.1) and very significant (**, p<0.01) differences are indicated in the cells.*

**Supplemental Table S2 (linked to Figure 2): The larval chemotactic behaviour is unaffected in homozygous Klp64D$^{kj}$ mutants.**

| *Stimulus* | Ethyl Acetate ($10^{-5}$) | | | Butanol ($10^{-2}$) | | | Fructose (1 M) | | |
|---|---|---|---|---|---|---|---|---|---|
| *Genotype* | Mean | S.D. | n | Mean | S.D. | n | Mean | S.D. | n |
| *Canton S* | 0.79 | 0.12 | 118 | 0.61 | 0.14 | 123 | 0.671 | 0.129 | 79 |
| *Kj353* | 0.83 | 0.08 | 32 | 0.69 | 0.13 | 32 | 0.671 | 0.093 | 24 |
| *Kj429* | 0.83 | 0.10 | 27 | 0.65 | 0.12 | 27 | 0.610 | 0.089 | 24 |
| *Kj925* | 0.80 | 0.10 | 35 | 0.63 | 0.13 | 28 | 0.669 | 0.120 | 26 |
| *Kj1072* | 0.83 | 0.11 | 52 | 0.55 | 0.13 | 51 | 0.466 | 0.113 | 37 |
| *Kj1070* | 0.88 | 0.07 | 30 | 0.69 | 0.12 | 32 | 0.678 | 0.096 | 25 |
| *OR83b$^{1}$* | 0.03 | 0.22 | 26 | 0.08 | 0.15 | 21 | 0.760 | 0.090 | 24 |

*Note: n-vale indicates total number of runs counted. Each run lasted for 15 minutes and consisted of approximately 20 larvae.*

**Supplemental Table S3: List of fly stocks used:**

| Fly stocks Used | Nature | Reference |
|---|---|---|
| *ChaTGal4* | Transgene, recombinant Gal4 under *Cha* promoter, expresses in cholinergic neurons | Salvaterra and Kitamoto, 2001 |
| *Or83bGal4* | Transgene, recombinant Gal4 under *Or83b* promoter, expresses in odour sensing neurons | Wang et al., 2003 |
| *UAS-eGFP* | Recombinant eGFP transgenes expressed under Gal4/UAS promoter | Spana, 1999.9.27, P[34] constructs and insertions from Eric Spana. |
| *UAS- GFP: Tub84B* | Transgene, N-terminal fusion of eGFP to tubulin84B, cloned under the UAS enhancer elements. | Avidor-Reiss et al., 2004 |
| *UAS-GFP:Orco* | Transgene, N terminal fusion of eGFP to Orco insert with ORF and 3'UTR corresponding to nucleotide 168-1917. | Benton et al.,2006 |
| *UAS-GFP:OR47b* | Transgene, N terminal fusion of eGFP to Or47b ORF and 3'UTR | Benton et al.,2006 |
| *Orco:GFP$^{fosmid}$* | Fosmid line expressing C terminal fusion of GFP to Orco, under its endogenous promoter. | Sarov et al., 2016 |
| *UAS-mCD8:GFP* | eGFP Fusion between mouse lymphocyte marker CD8 and the green fluorescence protein | Lee and Lou et al., 1999 |
| *UAS-mCD8:RFP* | Recombinant, mouse lymphocyte marker CD8 and the red fluorescence protein | Yang et al., 2008 |

| | | |
|---|---|---|
| *Jupiter:*GFP | Protein trap, GFP inserted in place of first intron of the endogenous Jupiter Protein. | Karpova et al., 2006 |
| *UAS-Klp64D* | Recombinant full-length KLP64D Transgene | Ray et al., 1999 |
| *UAS-Klp64D:GFP* | Recombinant full-length KLP64D Transgene with C-terminal GFP fusion | Jana et al, 2011 |
| *UAS-Klp68D:YFP* | Recombinant full-length KLP68D Transgene with C-terminal YFP fusion | Jana et al, 2011 |
| *Klp64D*$^{k1}$ | Amorphic allele, uncoordinated adults, missense mutation (G13*) | Ray et al., 1999 |
| *Klp64D*$^{kj353}$<br>*Klp64D*$^{kj925}$<br>*Klp64D*$^{kj1072}$ | Hypomorphic *Klp64D* allele; fails to complement the odour reception defects with lethal/uncoordinated, *Klp64D* alleles, such as *Klp64D*$^{k1}$, *Klp64D*$^{k5}$ and *Klp64D*$^{k4}$. | Anjusha K, Jana SC and Ray K, this manuscript. |
| Klp68D$^{EY00199}$ or Klp68D$^{P1}$ | Hypomorph, Recombinant P-P{EPgy2}<w+>element inserted in the 5'UTR. | Bellen et al., 2004 |
| *UAS-Oseg-2:GFP* | eGFP Fusion between *Drosophila* orthologue of the IFT-72, Oseg-2 | Avidor-Reiss et al., 2004 |
